# APE1 assembles biomolecular condensates to promote the ATR-Chk1 DNA damage response in nucleolus

**DOI:** 10.1101/2022.08.22.504787

**Authors:** Jia Li, Haichao Zhao, Anne McMahon, Shan Yan

## Abstract

Multifunctional protein APE1/APEX1/HAP1/Ref-1 (designated as APE1) plays important roles in nuclease-mediated DNA repair and redox regulation in transcription. However, it is unclear how APE1 regulates the DNA damage response (DDR) pathways and influences genome integrity directly or indirectly. Here we show that siRNA-mediated APE1-knockdown or APE1 inhibitor treatment attenuates the ATR-Chk1 DDR under stress conditions in multiple immortalized cell lines. Congruently, APE1 overexpression (APE1-OE) activates the ATR DDR under unperturbed conditions, which is independent of APE1 nuclease and redox functions. Structural and functional analysis reveals a direct requirement of the extreme N-terminal 33 amino acids (NT33) within APE1 in the assembly of distinct biomolecular condensates *in vitro* and DNA/RNA-independent activation of the ATR DDR. Overexpressed APE1 co-localizes with nucleolar NPM1 and assembles biomolecular condensates in nucleoli in cancer but not non-malignant cells, which recruits ATR and its direct activator molecules TopBP1 and ETAA1. APE1 W119R mutant is deficient in nucleolar condensation and liquid-liquid phase separation and is incapable of activating nucleolar ATR DDR. Lastly, APE1-OE-induced nucleolar ATR DDR activation leads to compromised ribosomal RNA transcription and reduced cell viability. Taken together, we propose distinct mechanisms by which APE1 regulates ATR DDR pathways and functions in genome integrity maintenance.

## INTRODUCTION

It is estimated that more than 10,000 Apurinic/Apyrimidinic (AP) sites are produced per day in each mammalian cell under unperturbed conditions (1–3). AP sites are typically repaired by base excision repair (BER) and nucleotide excision repair (NER) mechanisms in normal cells (4–6). In the BER pathway, AP sites are catalyzed into single strand breaks (SSBs) with heterogenous 3’ and/or 5’ termini via several mechanisms including hydrolysis catalyzed by AP endonuclease 1 (APE1) and β- or β,δ-elimination by bifunctional DNA glycosylases (3,7,8). SSBs are primarily repaired via a global PARP1/XRCC1-mediated SSB repair pathway into intact DNA, while other homology-based DNA repair mechanisms may serve as auxiliary(9–11). Delayed or improper repair of AP sites and SSBs compromises DNA replication and transcription programs, which can ultimately result in cancer and neurodegenerative disorders (9,12–14). To maintain genome integrity, the ATM-Chk2 DNA damage response (DDR) pathway is activated by DNA double-strand breaks (DSBs) through a dimer dissociation mechanism of ATM and the recruitment of the Mre11-Rad50-Nbs1 (MRN) complex (15–17). As a sensor of reactive oxygen species (ROS), ATM DDR is activated by oxidative stress via a cysteine-mediated ATM dimer formation (18). In addition, stalled DNA replication forks and assorted DNA damage events can activate the ATR-Chk1 DDR pathway (2,19,20), in which TopBP1 and ETAA1 play direct roles in the activation of ATR kinase via distinct but evolutionarily conserved ATR activation domain (AAD) (21–25). RPA-coated ssDNA (RPA-ssDNA) at damage site is widely accepted as a scaffolding for ATR-ATRIP recruitment and activation (20, 26). We have demonstrated recently that the ATR DDR pathway can also be activated by oxidative DNA damage and plasmid-based defined SSB structures in *Xenopus* egg extracts(27–29). It remains unknown whether and how AP sites and SSBs can trigger ATR-mediated DDR to maintain genome stability in mammalian cell systems.

Accumulating evidence has revealed that the ATM/ATR-mediated DDR pathways can be activated in the nucleolus, a non-membrane bound subnuclear compartment known for ribosomal RNA (rRNA) synthesis and organized by nucleolar organizing regions (NORs) located on acrosomal chromosomes (chromosome 13, 14, 15, 21, and 22 in humans)(30). Nucleoli are composed of three sub-nucleolar compartments: the fibrillar center (FC), the dense fibrillar component (DFC), and the granular component (GC) (31). It is generally accepted that pre-rRNA is transcribed from rDNA in the FC and/or DFC. FCs are enriched in components of the RNA Pol I machinery such as UBF, whereas the DFC harbors pre-rRNA processing factors such as FIB1. Both the FC and the DFC are enclosed by the GC, which is enriched in NPM1 for pre-ribosome subunit assembly (32). Ribosomal DNA (rDNA) in the nucleolus is transcribed into rRNA by RNA polymerase I, and such rDNA regions are often challenged by various DNA damage likely due to high rate of transcription (33). Previous work has revealed that in response to nucleolar DSBs, ATM is activated as a nucleolar DDR (nDDR) to inhibit RNA polymerase I-mediated rRNA transcription and to recruit homologous recombination (HR)-directed DNA repair machinery such as BRCA1, 53BP1, RPA32, and Rad51/Rad52 for faithful repair independent of cell cycle stages (34, 35). Mechanistic studies demonstrated the implication of Nbs1, Mdc1, and Treacle (also known as TCOF1) in the ATM-mediated inhibition of rRNA transcription in nDDR (34,36–39). Interestingly, site-specific DSB generation in nucleoli by CRISPR/Cas9 or I-PpoI endonuclease also triggers the activation of an ATR-dependent nDDR, which functions downstream of the ATM/Treacle/MRN-mediated nDDR; however, DSB-induced ATR nDDR does not include Chk1/Chk2-mediated cell cycle arrest (35,39,40). In contrast, hypoosmotic stress activates TopBP1/Treacle-dependent ATR nDDR in nucleoli, due to increased R-loop stabilization and RPA-ssDNA formation on actively transcribed rDNA, which functions upstream of ATM nDDR (41). Moreover, DNA replication stress induced by hydroxyurea or aphidicolin also triggers a Treacle/TopBP1-dependent ATR-mediated nDDR activation in nucleoli (42). TopBP1 overexpression can also induce ATR nDDR and suppression of rRNA transcription in the absence of rDNA damage breaks (43). Nevertheless, our understanding on the molecular mechanisms of ATR-mediated nDDR pathways remains incomplete.

The multifunctional protein APE1/APEX1/HAP1/Ref-1 (designated as APE1) exhibits AP endonuclease, 3’-5’ exonuclease, 3’-phosphodiesterase as well as 3’ RNA phosphatase and 3’ exoribonuclease activities and has been implicated in the BER pathway and redox regulation of transcription activation (44–47). APE1 is an essential gene for early embryonic development in mice and APE1-null cells do not generally survive (48, 49). APE1 protein exhibits subcellular localization in the nuclei and mitochondria which is mediated by its extreme N-terminal 33 amino acids (NT33) and its extreme C-terminal 30 amino acids (CT30), respectively (50–52). APE1 is also shown to associate with nucleolar protein Nucleiophosmin (NPM1) and participants in RNA quality control and RNA metabolism (53–55). Whereas the role of APE1 in the processing of AP sites via its endonuclease activity is widely accepted, APE1 exonuclease activity in genome integrity has not been recognized until recently (56–58). In particular, biochemical, biophysical, and structural model strongly supports the notion that APE1 preferentially senses and binds to SSB structures (56–58). We have demonstrated that APE1 plays an upstream role of APE2 in the SSB-induced ATR DDR pathway activation in *Xenopus* egg extracts (29,56,59). While APE2 is demonstrated as a general regulator of the global ATR DDR pathway in pancreatic cancer cells (60), it remains unclear whether APE1 plays direct roles in the activation of ATR-dependent nuclear and/or nucleolar DDR pathway in mammalian cells.

Liquid-liquid phase separation (LLPS) by biomolecular condensates has been recently reported in several cellular processes (61). At the molecular level, nucleoli in *Xenopus laevis* oocytes can form multiple LLPS that underlies nucleolar sub-compartments by NPM1 and fibrillarin (FIB1) (32,62,63). Inducible ectopic expression of TopBP1’s AAD domain is sufficient to activate a global ATR DDR but not ATR nDDR driving cells into p53-mediated senescence (64). Intriguingly, ectopic overexpression of full-length TopBP1 induces ATR nDDR under unperturbed conditions (43). These studies suggest that TopBP1 regions other than the AAD domain have the capability to trigger ATR nDDR. Interestingly, TopBP1 undergoes LLPS *in vitro* and assembles nuclear condensates to switch on ATR DDR signaling (65, 66). Although ∼49 APE1-interacting proteins (e.g., NPM1, FUS, APP, LGALS3, and HNRNPA1) are involved in biomolecular condensates (67, 68); however, it remains unknown whether and how APE1 forms LLPS.

Here, we provide evidence using cultured mammalian cells and reconstitution systems that APE1 plays critical functions in the ATR-Chk1 DDR signaling pathway via several distinct regulatory mechanisms. siRNA-mediated APE1-knockdown (APE1-KD) or APE1 nuclease specific inhibitors compromised the ATR-Chk1 DDR pathway under different stress conditions in several cell lines, suggesting that APE1 and its nuclease activity are important for the ATR-Chk1 DDR pathway. Overexpression of wild type (WT) or mutant APE1, that lacks nuclease or redox function, activated ATR-Chk1 DDR pathway in cultured cells under unperturbed conditions, suggesting that APE1 plays a previously unidentified role in ATR DDR activation via a catalytically independent fashion. Excess addition of recombinant APE1 protein in nuclear extracts directly activated the ATR-Chk1 DDR pathway, which required APE1 NT33 motif but not nuclease/redox functions. APE1 assembled distinct biomolecular condensates in a DNA/RNA-independent manner to associate and recruit ATR, TopBP1, and ETAA1 in nuclear extracts *in vitro*. APE1-OE co-localized with NPM1 and recruited ATR, TopBP1, and ETAA1 to nucleoli in cultured cells to activate ATR nDDR pathway. Elevated DNA damage load, cell cycle arrest, decreased rRNA transcription, and reduced cell viability underlie the physiological significance of the distinct ATR nDDR pathway via APE1-assembled biomolecular condensates in nucleolus. Taken together, our results shed new light on the distinct regulation of global and nucleolar ATR DDR pathway by APE1 in genome integrity maintenance.

## MATERIAL AND METHODS

### Cell culture and preparation of total cell lysates and nuclear extracts

MDA-MB-231, PANC1, and U2OS cells were purchased from ATCC, and cultured in Dulbecco’s Modified Eagle’s Medium (DMEM) supplemented with 10% FBS and penicillin (100 U/mL) and streptomycin (100 μg/mL) at 37°C in CO_2_ incubator (5%). For stress condition experiments, cells were treated with H_2_O_2_ (1.25 mM), Camptothecin (CPT, 10 μM), or methyl methanesulfonate (MMS, 0.3 mg/mL) and incubated for 2 hours before cell collection and further analysis. After different treatments, cells were washed with phosphate-buffered saline (PBS) followed by resuspension in Lysis Buffer A (20 mM Tris-HCL pH 8.0, 150 mM NaCl, 2 mM EDTA, 0.5% Nonidet P-40, 0.5 mM Na_3_V0_4_, 5 mM NaF, 5 μg/mL of Aprotinin and 10 μg/mL of Leupeptin). The total cell lysates were isolated by centrifugation at 13,000 rpm for 30 minutes at 4°C.

The nuclear extracts were prepared as previously described (56) and briefly as follows. After washing with PBS and resuspension in Solution A (20 mM Tris-HCL pH 7.4, 10 mM NaCl, 3 mM MgCl_2_), cells were incubated on ice for 15 minutes, supplemented with Nonidet P-40 (a final concentration of 0.5%), and vortexed for 10 seconds. The samples were centrifuged at 3,000 rpm for 10 minutes to separate permeabilized nuclei from cytoplasmic fraction. The recovered nuclei were lysed with Lysis Buffer A and centrifuged at 13,000 rpm at 4°C for 30 minutes to prepare nuclear extracts.

### APE1 overexpression and siRNA-mediated knockdown experiments in cells

For overexpression assays, various expression plasmids (e.g., YFP, WT/mutant YFP-APE1, tGFP, tGFP-APE1, or △N33 tGFP-APE1, etc.) was added to cells at about 30% confluence via Lipofectamine 2000 transfection method and cultured for different times as indicated. For tGFP and tGFP-tagged protein expression, different doses of doxycycline was added 1 day after plasmid transfection and continued culture for protein expression induction for another 2 days.

For siRNA-mediated knockdown experiments, various siRNA as indicated was added to cells at 30% confluence using Lipofectamine RNAiMAX reagent transfection method. A non-target siRNA was used as CTL siRNA (5’-UGGUUUACAUGUCGACUAA-3’). The nucleotide sequences of siRNA On-Targetplus SMARTpool for APE1 include 5’-CAAAGUUUCUUACGGCAUA-3’, 5’-GAGACCAAAUGUUCAGAGA-3’, 5’-CUUCGAGCCUGGAUUAAGA-3’, and 5’-UAACAGCAUAUGUACCCUAA-3’. The nucleotide sequences of siRNA On-Targetplus SMARTpool for for NPM1 are 5’-GUAGAAGACAUUAAAGCAA-3’, 5’-AAUGCAAGCAAGUAUAGAA-3’, 5’-ACAAGAAUCCUUCAAGAAA-3’, and 5’-UAAAGGCCGACAAAGAUUA-3’. The nucleotide sequences of siRNA On-Targetplus SMARTpool for TopBP1 are 5’-ACAAAUACAUGGCUGGUUA-3’, 5’-ACACUAAUCGGGAGUAUAA-3’, 5’-GAGCCGAACAUCCAGUUUA-3’, and 5’-CCACAGUAGUUGAGGCUAA-3’.

### Recombinant DNA and proteins

Plasmid pET28HIS-hAPE1 (for WT His-APE1 protein) was a gift from Primo Schaer (Addgene plasmid #70757; http://n2t.net/addgene:70757; RRID:Addgene_70757) (69). Plasmid pcDNA3-YFP (for YFP protein) was a gift from Doug Golenbock (Addgene plasmid #13033; http://n2t.net/addgene:13033; RRID:Addgene_13033). Plasmid pcDNA3-YFP-APE1 (for WT YFP-APE1 protein) was prepared by PCR full-length APE1 from pET28HIS-hAPE1 into pcDNA3-YFP at BamHI and EcoRI sites. Recombinant pET28HIS-APE1-YFP (for His-APE1-YFP protein) was prepared by PCR WT YFP-APE1 from plasmid pcDNA3-YFP-APE1 and subcloned into pET28HIS vector at BamHI and HindIII sites. Plasmid pCMV6-AC-GFP-rtTA-APE1 (for WT tGFP-APE1 protein) was prepared by PCR full-length APE1 from pET28HIS-hAPE1 and subcloned into pCMV6-AC-GFP-rtTA vector (Origene #PS100125) at AscI and RsrII sites. Various △N33 APE1 deletion plasmids and pET28HIS-NT33-APE1-YFP were also constructed using similar approach to WT plasmids. Various point mutant plasmids were prepared using QuikChange II XL site-directed mutagenesis kit (Agilent). QIAprep spin miniprep kit was utilized to make recombinant plasmids following vendor’s protocol. Various His-tagged recombinant proteins were expressed and purified in E. coli DE3/BL21. Purified recombinant proteins were examined and verified on SDS-PAGE gels with coomassie staining.

### FAM-labeled DNA and RNA structures

The 39-bp FAM-labeled dsDNA-AP structure and 70-bp FAM-labeled dsDNA structure have been described recently (56). The 39-bp FAM-labeled dsDNA-AP structure was prepared by annealing of two complementary oligos (Forward #1: [FAM]-5’-TGCTCGTCAAGAGTTCGTAA[THF]ATGCCTACACTGGAGATC -3’; Reverse #1: 5’-GATCTCCAGTGTAGGCATCTTACGAACTCTTGACGAGCA -3’) as previously described (56). The 70-bp FAM-labeled dsDNA structure was prepared by annealing two complementary oligos (Forward #2: [FAM]-5’-TCGGTACCCGGGGATCCTCTAGAGTCGACCTGCAGGCATGCAAGCTTGGCGTAATCATGGTCATAGCTGT -3’; Reverse #2: 5’-ACAGCTATGACCATGATTACGCCAAGCTTGCATGCCTGCAGGTCGACTCTAGAGGATCCCCGGGTACCGA -3’). The dsDNA structure was treated with Nt.BstNBI and CIP to make the 70-bp FAM-labeled dsDNA-SSB structure(56). The 25-bp FAM-labeled ssRNA was synthesized by IDT ([FAM]-5’-GCAGCUGGCACGACAGGUAUGAAUC -3’).

### Immunoblotting analysis and antibodies

Immunoblotting analysis was performed as described previously (27-29,56,60). Primary antibodies were purchased from respective vendors: APE1 (Santa Cruz Biotechnology Cat#sc-17774), ATR (Santa Cruz Biotechnology Cat#515173), ATRIP (Santa Cruz Biotechnology Cat#365383), Chk1 (Santa Cruz Biotechnology Cat#sc-8408), Chk1 phosphorylation at Ser345 (Cell Signaling Technology Cat#133D3), eGFP (Life Technologies Corporation Cat#TA150041), ETAA1 (Abcam Cat#ab122245), NPM1 (Santa Cruz Biotechnology Cat#sc-47725), p53 phosphorylation at Ser15 (Cell Signaling Technology Cat#9284), p53 (Santa Cruz Biotechnology Cat#sc-126), PCNA (Santa Cruz Biotechnology Cat#sc-56), RPA32 (Thermo Fisher Scientific Cat#MA1-26418), RPA32 phosphorylation at Ser33 (Bethyl Laboratories Cat#A300-246A), TopBP1 (Santa Cruz Biotechnology Cat#271043), Tubulin (Santa Cruz Biotechnology Cat#sc-8035), YFP (BioVision Cat#3991-100).

### In vitro pull-down assays

For the pull-down experiments, 20 μg of various His-tagged recombinant protein was added to 100 μL of cell nuclear extracts. After a 4h-incubation, an aliquot of the mixture was collected as Input and the remaining mixture was supplemented with 100 μL of Interaction Buffer (25 mM imidazole in PBS, pH 7.0) that contains 10 μL of Ni-NTA beads. After incubation with rotation for 1 hour at room temperature or overnight at 4°C, bead-bound fractions were washed twice with Interaction Buffer. The Input and Pulldown (i.e., bead-bound fractions) samples were examined via immunoblotting analysis as indicated.

### Immunofluorescence microscopy analysis

After 24-hour seeding on coverslide, cells were transfected with different expression plasmids for 72 hours or as indicated. After washing with PBS, Cells were then fixed in 3% paraformaldehyde within PBS for 15 minutes and incubated with PBST (PBS supplemented with 0.2% Triton X-100) for 5 minutes. After 1-hour incubation in blocking buffer (5% BSA in PBS), cells on coverslide were incubated with different antibodies conjugated with AF647 fluorescence in blocking buffer at 4°C overnight. After PBS wash three times, coverslips were stained with DAPI for 5 minutes and were mounted in ProLong Gold Antifade Mountant. Fluorescence images were acquired on a DeltaVision Elite Deconvolution microscope system equipped with a DV Elite CMOS camera and 60x objective and were further edited with Fiji-ImageJ software. Most antibodies-AF647 were purchased from Santa Cruz Biotechnology (NPM1: Cat#sc-47725 AF647; ATR: Cat#sc-515173 AF647; TopBP1: Cat#sc-271043). Anti-ETAA1 AF647 was prepared by conjugating AF647 to anti-ETAA1 antibodies (Abcam Cat#ab122245) using the Alexa Fluor 647 Conjugation Kit (fast)-Lighting-Link (Abcam Cat#ab269823) following vendor’s protocol.

### APE1 endo/exonuclease assays in vitro

For *in vitro* endonuclease assays, the dsDNA-AP structure was treated with different doses of purified recombinant WT/mutant His-APE1 with/without APE1 inhibitors in APE1 Reaction Buffer (60 mM NaCl, 2 mM MgCl_2_, 2 mM DTT, 50 mM HEPES pH 7.4) at 37°C. Endonuclease assay reaction was quenched with equal volume of TBE-Urea Sample Buffer and denatured for 5 minutes at 95°C. For *in vitro* exonuclease assays, the dsDNA-SSB structure was added with different concentrations of purified recombinant WT/mutant His-APE1 protein with/without APE1 specific inhibitors in APE1 Reaction Buffer at 37°C. Exonuclease assay reactions were quenched with equal volume of TBE-Urea Sample Buffer and denatured at 95°C for 5 min. After a quick spin, samples from endo/exonuclease assays were examined on 15% TBE-Urea PAGE gel and imaged with a Bio-Rad ChemiDoc MP Imaging System.

### APE1 liquid-liquid phase separation (LLPS) assays in vitro

APE1 phase separation in nuclear extracts was performed in reaction mixtures containing PANC1 nuclear extract (2 μg/μL) and different doses of WT/mutant His-APE1-YFP protein in an ATR activation buffer (10 mM HEPES-KOH pH 7.6, 50 mM KCl, 0.1 mM MgCl_2_, 1 mM PMSF, 0.5 mM DTT, 1 mM ATP), which was modified from a previously reported TopBP1 phase separation condition(65). APE1 phase separation in a buffer was performed in reaction mixture of different doses of WT/mutant His-APE1-YFP protein in a LLPS buffer (50 mM HEPES, pH 7.5, 20 mM KCl, 10 mM MgCl_2_ and 2 mM DTT). Reaction mixtures were incubated at 37°C for 15 minutes or otherwise as indicated, followed by immunofluorescence microscopy on PEG silanized slides. Images were captured on an inverted microscope using a 10x objective.

### Quantitative reverse-transcription PCR (qRT-PCR) assays

Total RNA was extracted from cells with Trizol reagent (Invitrogen Cat#15596026) according to vendor’s instructions. Purified RNA was treated with Dnase I to remove any possible DNA contamination. qRT-PCR assays were performed with SuperScript III Platinum SYBR Green One-Step qTR-PCR Kit (Invitrogen Cat#11736051) according to vendor’s standard procedure. qRT-PCT reactions were run in ABI 7500 FAST RT PCR system with parameters: 50°C for 5 minutes, 95°C for 5 minutes, followed by 40 cycles of PCR 95°C for 15 seconds and 60°C for 30 seconds. Primers targeting pre-r-RNA 5’ external transcribed region (5’-GGAAGGAGGTGGGTGGAC-3’ and 5’-GCGGTACGAGGAAACACCT-3’) and primers targeting Actin (5’-CTCTTCCAGCCTTCCTTCCT -3’ and 5’-AGCACTGTGTTGGCGTACAG-3’) were described recently (43). Pre-rRNA signals were normalized to Actin signals.

### FACS analysis, Comet assays, and MTT assays

For cell cycle profiling via FACS analysis, pelleted cells were washed by PBS and resuspended in 300 μL ice cold PBS. Then cells were mixed with 700 μL ice cold 100% ethanol and incubated at 4°C for at least 2 hours. Cells were washed by PBS again and treated with RNase within PBS for 15 minutes at 37°C, followed by resuspension in Staining Buffer (100 mM Tris-HCl, pH7.4, 150 mM NaCl, 1 mM CaCl_2_, 0.5 mM MgCl_2_, 0.1% NP-40, 5μg/mL DAPI) for FACS analysis. Comet assays were performed using the OxiSelect Comet Assay Kit (Cell Biolabs Cat#STA-351-5) with alkaline (pH > 7.0) or neutral condition (pH = 7.0) following vendor’s standard protocol. Cell nuclei were imaged using a fluorescence microscope with a FITC filter and DP Controller software. Images were analyzed using Comet Assay IV Lite software. For cell viability analysis using MTT (Thiazolyl blue tetrazolium bromide) assays, cultured cells in 96-well plate were examined using a procedure as recently described (60).

### Quantification, statistical analysis and reproducibility

The data presented are representative of three biological replicates unless otherwise specified. All statistics were performed using GraphPad Prism version 8 for Windows. Statistical significance was ascertained between individual samples using a parametric unpaired t-test. Significance is denoted by asterisks in each figure: *P<0.05; **P<0.01; ***P<0.001; ****P<0.0001; ns, no significance. Error bars represent the standard deviation (SD) for three independent experiments, unless otherwise indicated.

## RESULTS

### APE1 is important for the ATR-Chk1 DDR pathway activation under stress conditions in mammalian cells

Our recent study has shown that APE1 plays an essential role in the ATR-Chk1 DDR pathway activation in *Xenopus* HSS system (56). To explore the function of APE1 in DDR pathway in mammalian cells, we first found that H_2_O_2_-induced oxidative stress triggered Chk1 phosphorylation at Ser345 in human breast cancer cell line MDA-MB-231, suggesting the activation of the ATR DDR pathway (Figure 1A). Notably, the H_2_O_2_-induced Chk1 phosphorylation was significantly reduced in siRNA-mediated APE1-KD in MDA-MB-231 cells (Figure 1A). We observed a similar reduction of H_2_O_2_-induced Chk1 phosphorylation in human pancreatic cancer cell line PANC1 cells after APE1-KD (Figure 1D).

**Figure 1.**
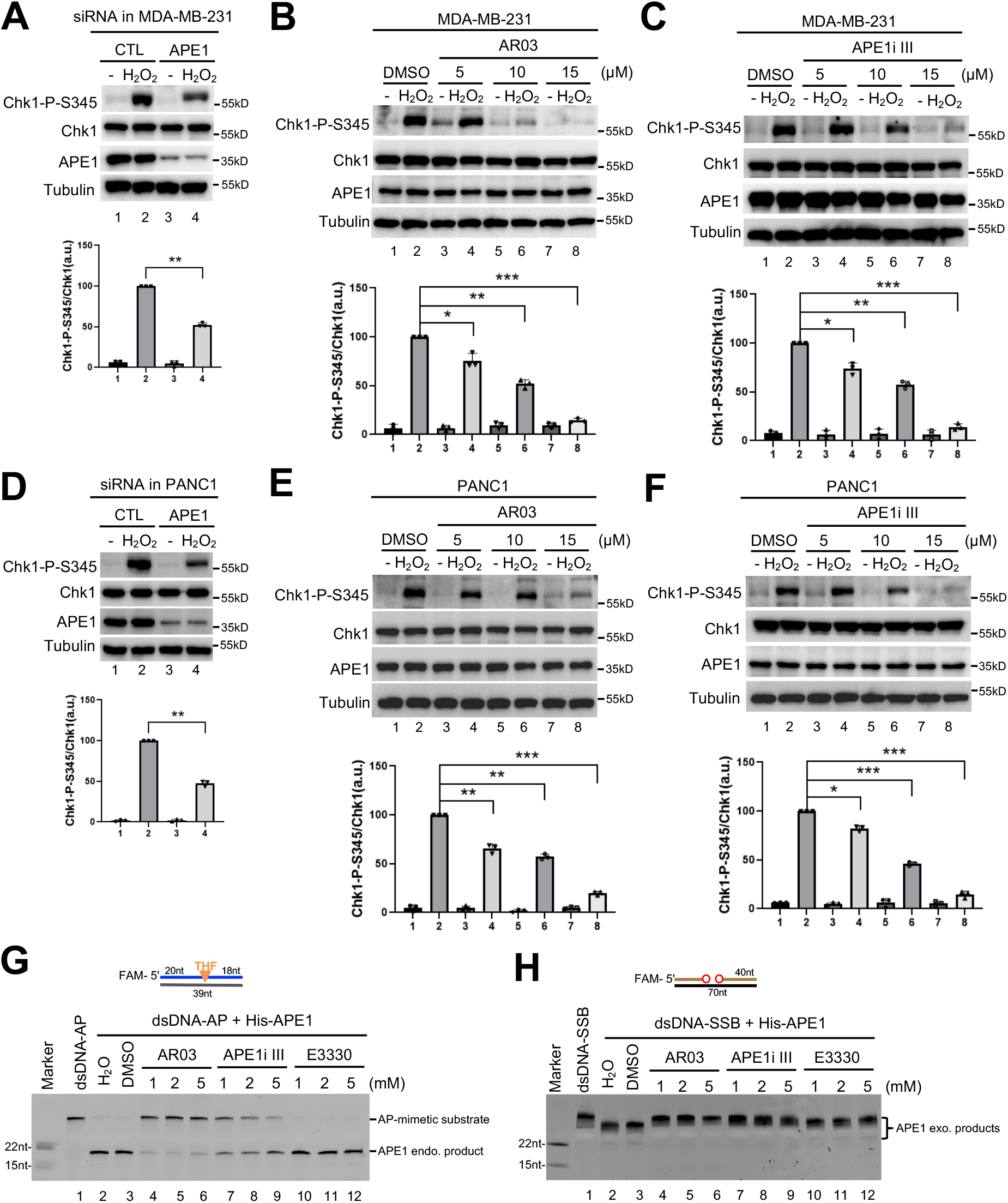
APE1 and its nuclease activity is important for ATR-Chk1 DDR pathway in mammalian cells. (**A**) siRNA-mediated APE1-knockdown impaired H_2_O_2_-induced Chk1 phosphorylation at Ser345 (Chk1-P-S345) in MDA-MB-231 cells. Total cell lysates were extracted and analyzed via immunoblotting analysis as indicated. “Chk1-P/Chk1” was quantified by the intensity of Chk1-P-S345 vs. total Chk1. Means ± SD, n=3. (**B-C**) The H_2_O_2_-induced Chk1 phosphorylation was compromised by AR03 or APE1i III at different doses (5, 10, or 15 μM) in MDA-MB-231 cells. Means ± SD, n=3. (**D**) The H_2_O_2_-induced Chk1 phosphorylation was reduced in APE1-KD PANC1 cells. Means ± SD, n=3. (**E-F**) The H_2_O_2_-induced Chk1 phosphorylation was impaired by AR03 or APE1i III in a dose-dependent manner (5, 10, or 15 μM) in MDA-MB-231 cells. Means ± SD, n=3. (**G**) AR03 or APE1i III but not E3330 inhibited APE1’s endonuclease activity. Different doses of AR03, APE1i III, or E3330 (1, 2, or 5 mM) was added to endonuclease assays containing 0.185 μM APE1 protein and a 5’-FAM-labeled dsDNA-AP substrate. After incubation for 30 minutes, samples were analyzed via 20% TBE gel. (**H**) AR03 or Inhibitor III could inhibit APE1’s exo-nuclease activity. Different doses of AR03, APE1i III, or E3330 (1, 2, or 5 mM) was added to exonuclease assays containing 0.185 μM APE1 protein and 5’-FAM-labeled dsDNA-SSB substrate. After 30-minutes incubation, samples were examined via 20% TBE gel.

Previous studies have identified and characterized AR03 and APE1 inhibitor III (APE1i III) as specific inhibitors for APE1 endonuclease activity and E3330 as an inhibitor of APE1 redox function (70–72). As expected, recombinant human full-length WT APE1 displayed AP endonuclease activity and 3’-5’ exonuclease activity in a dose-dependent manner in *in vitro* reconstitution systems (Supplementary Figure S1A-C). Importantly, H_2_O_2_-induced Chk1 phosphorylation was significantly reduced with the addition of AR03 or APE1i III in a dose-dependent manner in MDA-MB-231 cells (Figure 1B-C) and PANC1 cells (Figure 1E-F). However, the addition of E3330 had almost no noticeable effect on the H_2_O_2_-induced Chk1 phosphorylation in PANC1 cells (Supplementary Figure S1D). We verified that AR03 and APE1i III inhibited both AP endonuclease and 3’-5’ exonuclease activity of recombinant APE1 *in vitro*, whereas E3330 had minimal effects on APE1 nuclease activities (Figure 1G-H). These observations suggest that APE1 and its nuclease activity but not redox regulation is important for APE1’s function in oxidative stress-induced ATR-Chk1 DDR pathway in mammalian cells.

Furthermore, methyl methanesulfonate (MMS)-induced DNA alkylation damage and camptothecin (CPT)-induced DSBs and SSBs triggered Chk1 phosphorylation in MDA-MB-231 cells (Supplementary Figure S1E-H). MMS/CPT-induced Chk1 phosphorylation was also compromised by the addition of AR03 or APE1i III in a dose-dependent manner in MDA-MB-231 cells (Supplementary Figure S1E-H). AR03 and APE1i III similarly impaired MMS/CPT-induced Chk1 phosphorylation in PANC1 cells (Supplementary Figure S1I-L). Based on these evidences, APE1 may play a key role in the regulation of the ATR-Chk1 DDR pathway under stress conditions in mammalian cells.

### APE1 overexpression triggers ATR-Chk1 DDR pathway activation under unperturbed conditions in mammalian cells

Meta-analysis has revealed APE1 upregulation in cancer tissues compared with normal tissues and the association of APE1 overexpression with poor survival in patients with solid tumors (73, 74). To elucidate the potential role of APE1 overexpression in DDR pathway, we over-expressed YFP-tagged APE1 (YFP-APE1) to a similar level of endogenous APE1 in MDA-MB-231 cells, and found that YFP-APE1 but not YFP transfection nor control (CTL) transfection triggered Chk1 phosphorylation after 4 days of transfection (Figure 2A). Of note, the YFP-APE1-induced Chk1 phosphorylation almost disappeared 7 days after transfection (Figure 2A). Pre-treatment of ATR inhibitor VE-822 but not ATM inhibitor KU55933 nor DNA-PKcs inhibitor NU7441 compromised the YFP-APE1 overexpression-induced Chk1 phosphorylation in MDA-MB-231 cells (Figure 2B). This suggests that APE1 overexpression leads to ATR-dependent Chk1 phosphorylation and DDR pathway activation in cells under unperturbed conditions.

**Figure 2.**
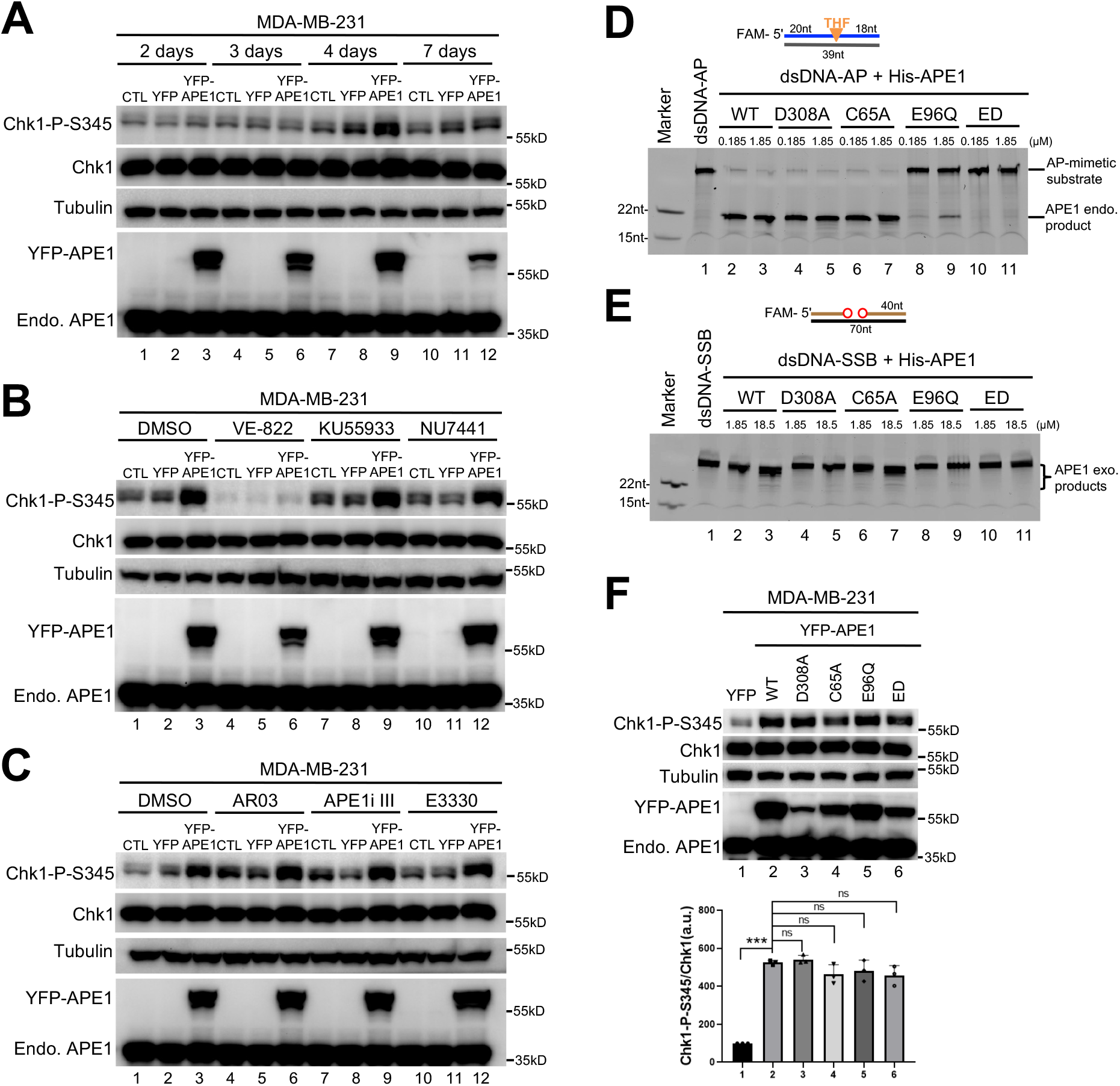
APE1 overexpression leads to the activation of ATR-Chk1 DDR pathway independent of its nuclease and redox function in mammalian cells. (**A**) Chk1 phosphorylation was activated by the overexpression of YFP-APE1 but not YFP nor CTL transfection (no vector transfection) in MDA-MB-231 cells. After different times of incubation (i.e., 2, 3, 4, or 7 days), total cell lysates were extracted and examined via immunoblotting analysis as indicated. (**B**) Chk1 phosphorylation induced by YFP-APE1 was dependent on ATR, but not ATM nor DNA-PKcs. After 4-day overexpression of YFP-APE1 or YFP, different DDR kinase inhibitors (5 μM of VE-822, KU55933, or NU7441) were added to MDA-MB-231 cells for 2 hours, followed by total cell lysate extraction and immunoblotting analysis as indicated. (**C**) APE1 inhibitors almost had no noticeable effect on Chk1 phosphorylation by YFP-APE1 overexpression. After 2-day overexpression of YFP-APE1 or YFP, different APE1 inhibitors (1 μM of AR03, APE1i III, or E3330) were added to MDA-MB-231 cells for 2 days, followed by total cell lysate extraction and immunoblotting analysis as indicated. (**D**) The endonuclease activity of WT or mutant (i.e., D308A, C64A, E96Q, ED) His-APE1 was examined on 20% TBE gel. (**E**) The exonuclease activity of WT or mutant (i.e., D308A, C64A, E96Q, ED) His-APE1 was examined on 20% TBE gel. (**F)** Chk1 phosphorylation was triggered by WT/mutant YFP-APE1 (i.e., D308A, C64A, E96Q, ED) overexpression in MDA-MB-231 cells. After 4-day overexpression of WT/mutant YFP-APE1 or YFP in MDA-MB-231, total cell lysates were extracted and analyzed via immunoblotting analysis. The data are presented as Means ± SD, n=3.

Next, we attempted to determine whether the nuclease activities and/or redox function of APE1 play important roles for the APE1 overexpression-induced ATR DDR pathway activation. First, we utilized various APE1 inhibitors and found, surprisingly, that none of the AR03, APE1i, and E3330 had noticeable effect on the APE1 overexpression-induced ATR DDR pathway (Figure 2C). Next, we constructed various mutants of His-tagged recombinant human APE1 (D308A, E96Q, E96Q/D210N (ED), and C65A) (75–78) and verified these mutants for APE1 nuclease activity *in vitro*. C65A His-APE1 had no noticeable deficiency in 3’-5’ exonuclease activity and AP endonuclease activity compared with WT His-APE1 (Figure 2D-E and Supplementary Figure S1A). While D308A His-APE1 was deficient for 3’-5’ exonuclease activity and proficient in AP endonuclease activity, E96Q and ED His-APE1 were defective for both 3’-5’ exonuclease activity and AP endonuclease activity (Figure 2D-E, and Supplementary Figure S1A). We then overexpressed various mutant YFP-APE1 in MDA-MB-231 cells for 4 days and found that all mutant YFP-APE1 (D308A, E96Q, ED, and C65A) still triggered Chk1 phosphorylation in a similar fashion as the WT YFP-APE1 (Figure 2F). Although we added the same amount of plasmid of WT/mutant YFP-APE1 for transfection, WT and mutant YFP-APE1 overexpression in MDA-MB-231 was slightly different between each other. Nonetheless, immunoblotting analysis and fluorescence microscopy analysis showed that all WT/mutant YFP-APE1 were indeed overexpressed as compared to endogenous APE1 levels (Figure 2F and Supplementary Figure S2A). These observations suggest that the APE1 overexpression-induced activation of the ATR-Chk1 DDR pathway under unperturbed conditions in MDA-MB-231 cells is independent of APE1 nuclease activities and redox function.

To diversity our results, we also overexpressed WT/mutant YFP-APE1 in PANC1 cells (Supplementary Figure S2B and S2D) and human osteosarcoma U2OS cells (Supplementary Figure S2C and S2E). We found that Chk1 phosphorylation was upregulated by overexpression of WT YFP-APE1 to slightly different levels in different cell lines (i.e., ∼5 folds in MDA-MB-231, ∼3.5 folds in PANC1, and ∼2.5 folds in U2OS) (Figure 2F and Supplementary Figure S2D-E). Similar to WT YFP-APE1, mutant YFP-APE1 (D308A, E96Q, ED, and C65A) also triggered Chk1 phosphorylation under unperturbed conditions in PANC1 cells and U2OS cells (Supplementary Figure S2B-E1). These results suggest that APE1 overexpression-induced ATR-Chk1 DDR pathway under unperturbed conditions is a conserved response in cultured mammalian cell lines.

### The NT33 motif of APE1 associates with ATR, ATRIP, and RPA to regulate the ATR-Chk1 DDR pathway in cells and in nuclear extracts

To test whether the different level of APE1-OE correlates with the level of ATR DDR activation, we constructed a doxycycline-inducible plasmid that induced overexpression of turbo-GFP (tGFP)-tagged APE1 (tGFP-APE1) at different levels. Overexpression of tGFP-APE1 but not tGFP at a concentration of 50 μg/mL but not 10 μg/mL doxycycline for three days induced Chk1 phosphorylation in PANC1 cells under unperturbed conditions (Figure 3A). This result suggests that the level of overexpression APE1 is important for Chk1 phosphorylation. Previous studies have shown that the extreme NT33 motif of human APE1 interacts with RNA and NPM1 (53,55,79). Interestingly, we found that overexpression of △NT33 tGFP-APE1 lacking the NT33 motif was deficient for Chk1 phosphorylation, although △NT33 tGFP-APE1 was expressed to a comparable level to WT tGFP-APE1 (Figure 3A). This observation suggests that the NT33 motif within APE1 is critical for the APE1-OE-induced ATR DDR activation. Prior research has identified a nuclear localization signal (NLS) within the NT33 motif of APE1 (51, 55). Our fluorescence microscopy analysis shows that similar to tGFP, △NT33 tGFP-APE1 does not localize to specific organelles, while WT tGFP-APE1 mostly localizes to nuclei (Figure 3B). This defective translocation of △NT33 tGFP-APE1 into nuclei may be one mechanism for its deficiency in ATR DDR pathway under unperturbed conditions.

**Figure 3.**
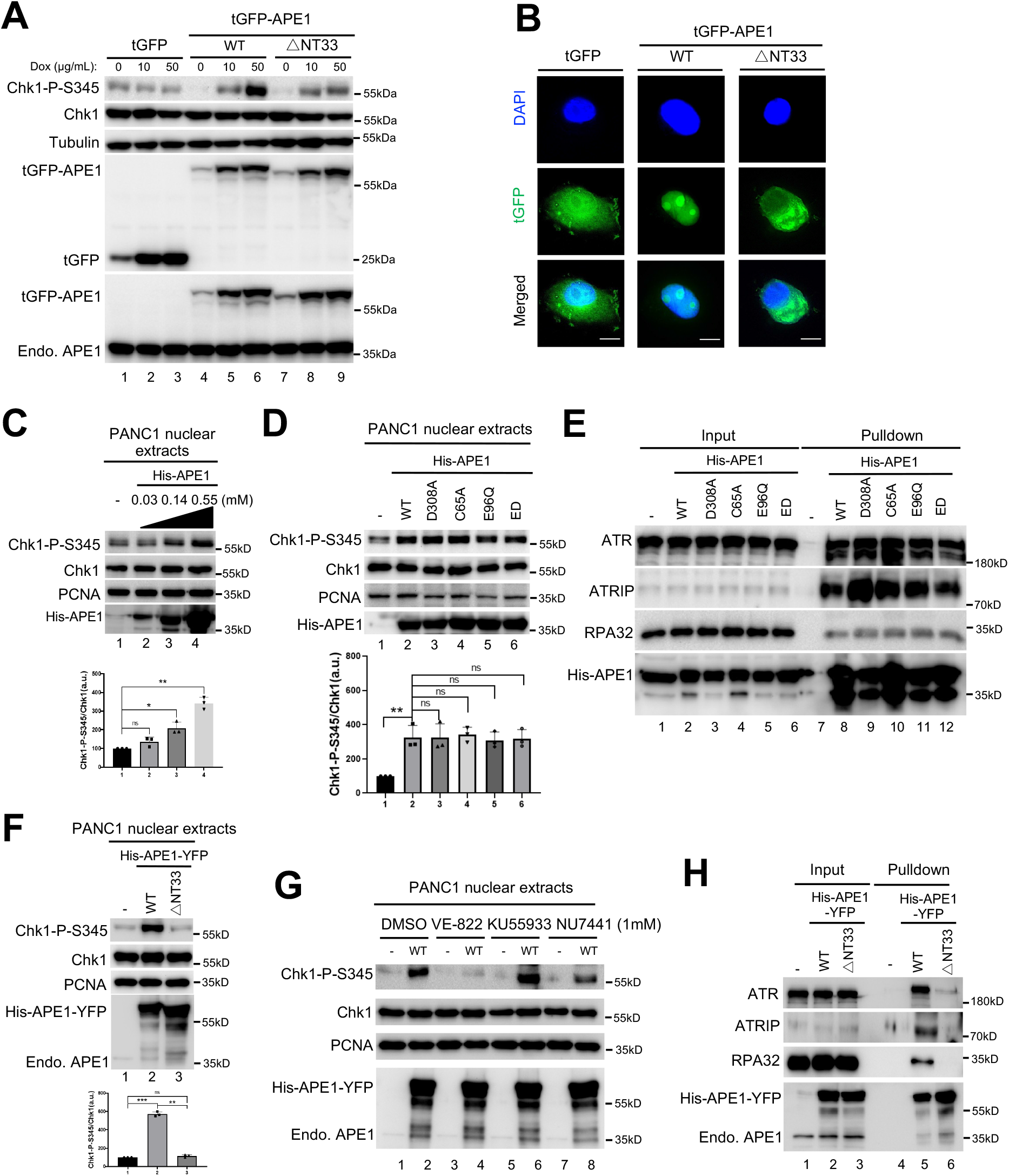
The APE1 N33 motif is required for ATR-Chk1 DDR pathway *in vivo* and *in vitro*. (**A**) Overexpression of WT but not △N33 tGFP-APE1 nor tGFP activated Chk1 phosphorylation in PANC1 cells. APE1 or N33-depletion over-expression plasmid was added to PANC1 cells. After 1-day incubation of various overexpression plasmid transfection, different doses of doxycycline (0, 10, or 50 μg/mL) were added to PANC1 cells for another 2 days. Total cell lysates were then extracted and analyzed via immunoblotting analysis as indicated. (**B**) △N33 tGFP-APE1 was deficient in nuclear localization in PANC1 cells. tGFP, WT or △N33 tGFP-APE1 overexpression plasmid was added to PANC1 cells. After 1-day incubation of overexpression plasmid transfection, 50 μg/mL doxycycline was added to PANC1 cells for another 2 days, followed with fluorescence microscopy analysis. Scale bar = 10 μm. (**C**) Chk1 phosphorylation was triggered by the excess addition of recombinant His-APE1 protein in a dose-dependent manner in PANC1 nuclear extracts. Different doses of His-APE1 protein were added to PANC1 nuclear extracts for incubation for 30min at room temperature. The samples were analyzed via immunoblotting analysis as indicated. Chk1-P/Chk1 was quantified and analyzed as means ± SD (n=3). (**D**) Chk1 phosphorylation induced by excess addition of recombinant His-APE1 protein in PANC1 nuclear extracts was independent of its nuclease activity and redox function. WT or mutant (i.e., D308A, C65A, E96Q, ED) His-APE1 was added to PANC1 nuclear extracts for 30-minutes incubation at room temperature, followed by immunoblotting analysis. Chk1-P/Chk1 was quantified and analyzed as means ± SD (n=3). (**E**) Pulldown assays suggest that WT/mutant His-APE1 associated with ATR, ATRIP and RPA32 in PANC1 nuclear extracts. “Input” or “Pulldown” samples were examined via immunoblotting analysis as indicated. (**F**) Chk1 phosphorylation was activated by excess addition of WT His-APE1-YFP protein but not △N33 His-APE1-YFP in PANC1 nuclear extracts. Chk1-P/Chk1 was quantified and analyzed as means ± SD (n=3). (**G**) Chk1 phosphorylation induced by WT His-APE1-YFP protein in PANC1 nuclear extracts was inhibited by VE-822 but not KU55933 nor NU7441 (1 mM). (**H**) Pulldown assays suggest that WT but not △N33 His-APE1-APE1 associated with ATR, ATRIP and RPA32 in PANC1 nuclear extracts. “Input” or “Pulldown” samples were examined via immunoblotting analysis as indicated.

To directly test the role of NT33 motif of APE1 in the ATR DDR activation but not via its role in nuclear localization, we established an *in vitro* ATR DDR activation system using nuclear extracts isolated from PANC1 cells and found that the addition of purified WT His-tagged APE1 (His-APE1) protein upregulated Chk1 phosphorylation significantly in a dose-dependent manner (Figure 3C). Similar to WT His-APE1, the addition of mutant His-APE1 (D308A, C65A, E96Q, and ED) also upregulated Chk1 phosphorylation in nuclear extracts (Figure 3D), consistent with the upregulation of Chk1 phosphorylation by the overexpression of WT/mutant YFP-APE1 in mammalian cells (Figure 2 and Supplementary Figure S2). We observed similar Chk1 phosphorylation through the addition of WT/mutant His-APE1 protein in nuclear extracts from U2OS cells (Supplementary Figure S3A-B). GST pulldown experiments showed that WT and mutant His-APE1 protein could pulldown ATR, ATRIP, and RPA32 in nuclear extracts from PANC1 cells (Figure 3E) and from U2OS cells (Supplementary Figure S3C). Furthermore, we expressed and purified recombinant WT and △NT33 APE1 with both His-tag and YFP-tag (His-APE1-YFP) for subsequent biochemical and imaging analyses. Of note, excess addition of WT His-APE1-YFP but not △NT33 His-APE1-YFP increased Chk1 phosphorylation significantly in nuclear extracts, although both recombinant proteins were added to comparable levels (Figure 3F). Moreover, the Chk1 phosphorylation induced by excess WT His-APE1-YFP in nuclear extracts was compromised by VE-822 but not KU55933 nor NU7441 (Figure 3G). Because there is no intact nuclear membrane in nuclear extracts, our observation provides the first evidence for the direct requirement of the APE1 NT33 motif in ATR DDR activation. Furthermore, GST pulldown experiments revealed that WT His-APE1-YFP but not △NT33 His-APE1-YFP associated with ATR, ATRIP, and RPA32 in PANC1 nuclease extracts (Figure 3H), suggesting that the NT33 motif of APE1 is important for APE1 to interact with ATR and its associated proteins for ATR activation in nuclear extracts. The addition of NT33 motif only to nuclear extracts did not trigger Chk1 phosphorylation (Supplementary Figure S3D), suggesting that NT33 motif within APE1 is required but not sufficient for Chk1 phosphorylation in nuclear extracts. These data suggest that nuclear localization and interaction with ATR and association proteins are two critical functions of the APE1 NT33 motif in the ATR DDR activation.

### APE1 assembles biomolecular condensates to recruit ATR, TopBP1, and ETAA for ATR DDR activation in nuclear extracts and formed LLPS in vitro

We next tested whether APE1 could form biomolecular condensates *in vitro*. Fluorescent microscopy analysis showed that recombinant WT His-APE1-YFP formed micrometer-sized biomolecular condensates in nuclear extracts in a dose and time-dependent manner (Figure 4A and Supplementary Figure S4A-B). In contrast, recombinant △NT33 His-APE1-YFP was deficient in the assembly of biomolecular condensates in nuclear extracts (Figure 4A and Supplementary Figure S4B). To test whether such APE1-induced condensates were dependent on DNA and/or RNA in nuclear extracts, we added DNase I and RNase A in nuclear extracts and found no noticeable change on the assembly of condensates by WT His-APE1-YFP (Figure 4B). None of APE1 specific inhibitors (AR03, APE1i III, and E3330) had noticeable effects on APE1-induced biomolecular condensates (Figure 4C) and Chk1 phosphorylation status in nuclear extracts (Figure 4D). Notably, pulldown assays showed that WT but not △NT33 His-APE1-YFP formed protein complexes with ATR and its activators TopBP1 and ETAA1 in nuclear extracts, for which treatment of AR03 or E3330 had no effect on the complex formation of WT His-APE1-YFP and ATR/TopBP1/ETAA1 (Figure 4E). These results suggest that WT but not △NT33 APE1 forms biomolecular condensates which in turn recruit ATR and its activators for ATR activation in nuclear extracts.

**Figure 4.**
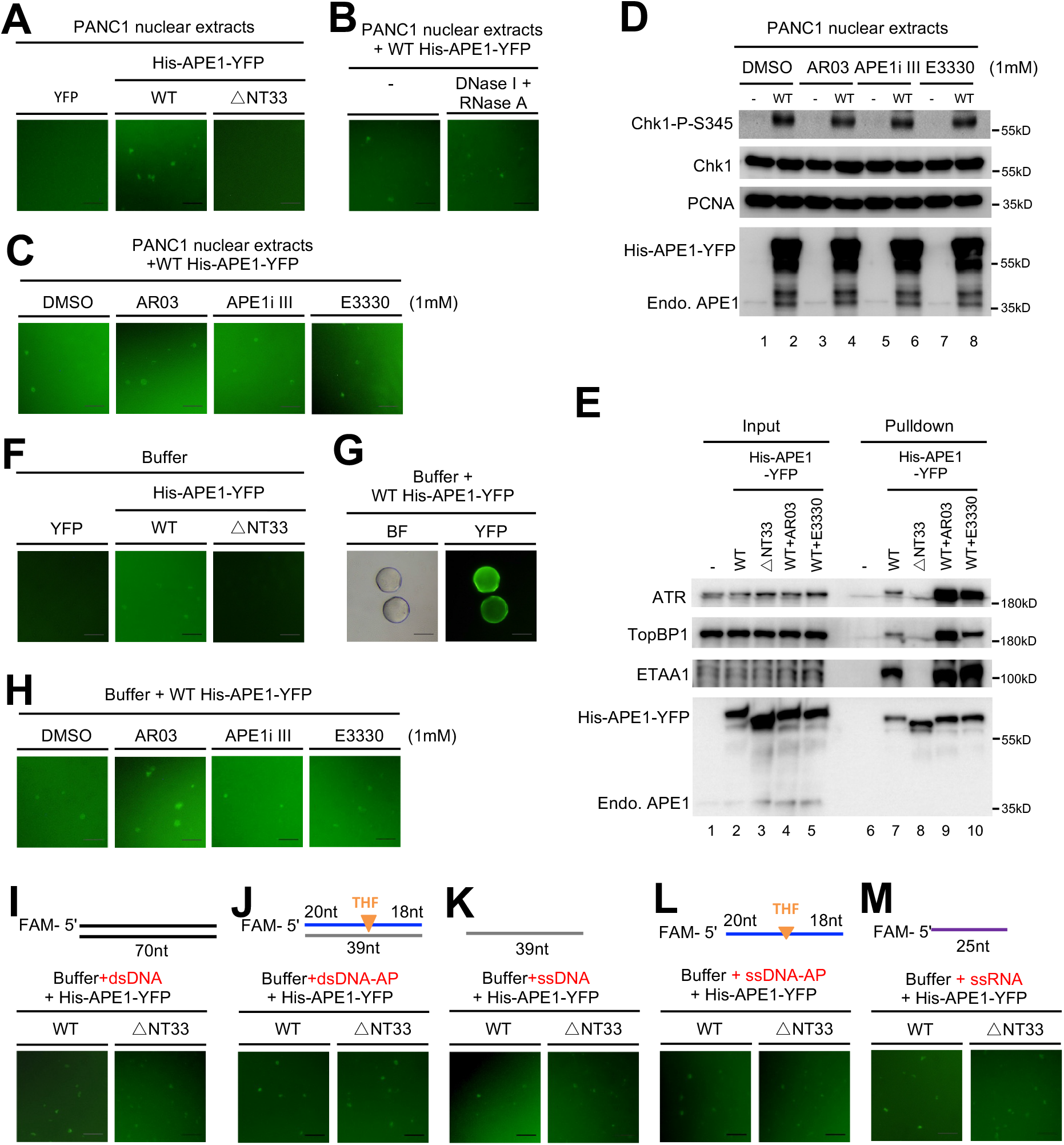
APE1 assembles biomolecular condensates *in vitro* to promote ATR DDR pathway. (**A**) WT but not △N33 His-APE1-YFP formed phase separation in PANC1 nuclear extracts. Recombinant YFP, WT or △N33 His-APE1-YFP protein (0.55 mM) was added to PANC1 nuclear extract and incubated for 15 minutes at room temperature, followed by fluorescence microscopy analysis. Scale bar = 50 μm. (**B**) DNase I and RNase A had almost no effect on the APE1-assembled phase separation in nuclear extracts. DNase I and RNase A (2 mg/mL each) was added to PANC1 nuclear extracts and incubated for 30 minutes at 37°C. Then WT His-APE1-YFP protein (0.55 mM) was added and incubated for 10 minutes at room temperature, followed by fluorescence microscopy analysis. Scale bar = 50 μm. (**C**) APE1 inhibitors had no noticeable effect on phase separation formed by APE1 in nuclear extracts. WT His-APE1-YFP protein (0.55 mM) was added to PANC1 supplemented with DMSO or APE1 inhibitors (AR03, APE1i III, and E3330). After 15-minute incubation at room temperature, reaction mixtures were examined via fluorescence microscopy analysis. Scale bar = 50 μm. (**D**) APE1 inhibitors had no effect on the Chk1 phosphorylation induced by the excess addition of APE1 protein in PANC1 nuclear extracts. APE1 inhibitor (AR03, APE1i III, or E3330) was added to PANC1 nuclear extracts and incubated for 10 minutes at room temperature, which was supplemented with WT His-APE1-YFP (0.55 mM) and incubated for 30 minutes at room temperature. The samples were then analyzed via immunoblotting analysis as indicated. (**E**) WT but not △N33 His-APE1-YFP associated with ATR, TopBP1 and ETAA1 in PANC1 nuclear extracts. “Input” and “Pulldown” samples from pulldown experiments were examined via immunoblotting analysis as indicated. AR03 or E3330 was added to 1mM. (**F**) WT but not △N33 His-APE1-YFP formed phase separation in a LLPS buffer. WT but not △N33 His-APE1-YFP (0.55 mM) was added to a buffer and incubated for 15 minutes at room temperature, followed by fluorescence microscopy analysis. Scale bar = 50μm. (**G**) Phase separation formed by recombinant WT His-APE1-YFP protein was fused into large biomolecular condensates in a LLPS buffer. WT His-APE1-YFP (0.55 mM) was added to a buffer and incubated for 2 hours before fluorescence microscopy analysis. BF, bright field. Scale bar = 50 μm. (**H**) Phase separation assembled by APE1 in a LLPS buffer was not affected by the APE1 inhibitors. WT His-APE1-YFP protein (0.55 mM) was added to PANC1 supplemented with DMSO or APE1 inhibitors (AR03, APE1i III, and E3330). After 15-minute incubation at room temperature, reaction mixtures were examined via fluorescence microscopy analysis. Scale bar = 50 μm. (**I-M**) The requirement of NT33 motif within APE1 for phase separation in a LLPS buffer was bypassed with the presence of different DNA or RNA. WT but not △N33 His-APE1-YFP (0.55 mM) was added to a buffer supplemented with 0.5 μM of different DNA/RNA structures (dsDNA (**I**), dsDNA with AP site (**J**), ssDNA (**K**), ssDNA with AP site (**L**) and ssRNA (**M**)). Reaction mixtures were incubated for 15 minutes at room temperature before fluorescence microscopy analysis. Scale bar = 50 μm.

To further explore whether APE1 can form LLPS *in vitro*, we found that recombinant WT His-APE1-YFP but not △NT33 His-APE1-YFP assembled biomolecular condensates in a dose and time-dependent manner in a LLPS buffer (Figure 4F and Supplementary Figure S4C-D). It should be noted that smaller condensates fused into larger circular condensates (Figure 4G and Supplementary Figure S4D), supporting the LLPS feature of APE1 protein *in vitro*. APE1-induced biomolecular condensates in the LLPS buffer are not sensitive to APE1 inhibitors (Figure 4H). Interestingly, although DNA/RNA is dispensable for the APE1-induced assembly of biomolecular condensates, the addition of different DNA/RNA structures (dsDNA, dsDNA with AP site, ssDNA, ssDNA with AP site, and ssRNA) bypassed the requirement of APE1 NT33 motif for condensate formation *in vitro* (Figure 4I-L).

### Overexpressed APE1 colocalized with NPM1 and assembles LLPS in vivo to recruit ATR and its activator proteins for nDDR in nucleoli in cancer cells but not non-malignant cells

As WT tGFP-APE1, but not △NT33 tGFP-APE1 nor tGFP, was found concentrated to some regions within nuclei (Figure 3B), we next asked whether and how APE1 forms biomolecular condensates within cell’s nuclei *in vivo*. First, our fluorescent microscopy analysis showed co-localization of nucleolar NPM1 with concentrated YFP-APE1 but not YFP in PANC1 cells, suggesting that overexpressed YFP-APE1 protein is translocated to nucleoli within nuclei (Figure 5A). We found similar observation in other two cancer cells MDA-MB-231 and U2OS cells (Supplementary Figure S5A-B). Furthermore, ATR and its activators TopBP1 and ETAA1 were all co-localized with YFP-APE1 but not YFP in nucleoli in PANC1 cells (Figure 5B-D). Intriguingly, overexpressed YFP-APE1 did not form condensates in nucleoli nor activated Chk1 phosphorylation in non-malignant human pancreatic duct epithelial (HPDE) cells (Figure 5E and Supplementary Figure S5C). These observations suggest that overexpressed YFP-APE1 assembles biomolecular condensates within nucleoli and recruits ATR, TopBP1, and ETAA1 for ATR nDDR activation in cancer cells but not non-malignant cells.

**Figure 5.**
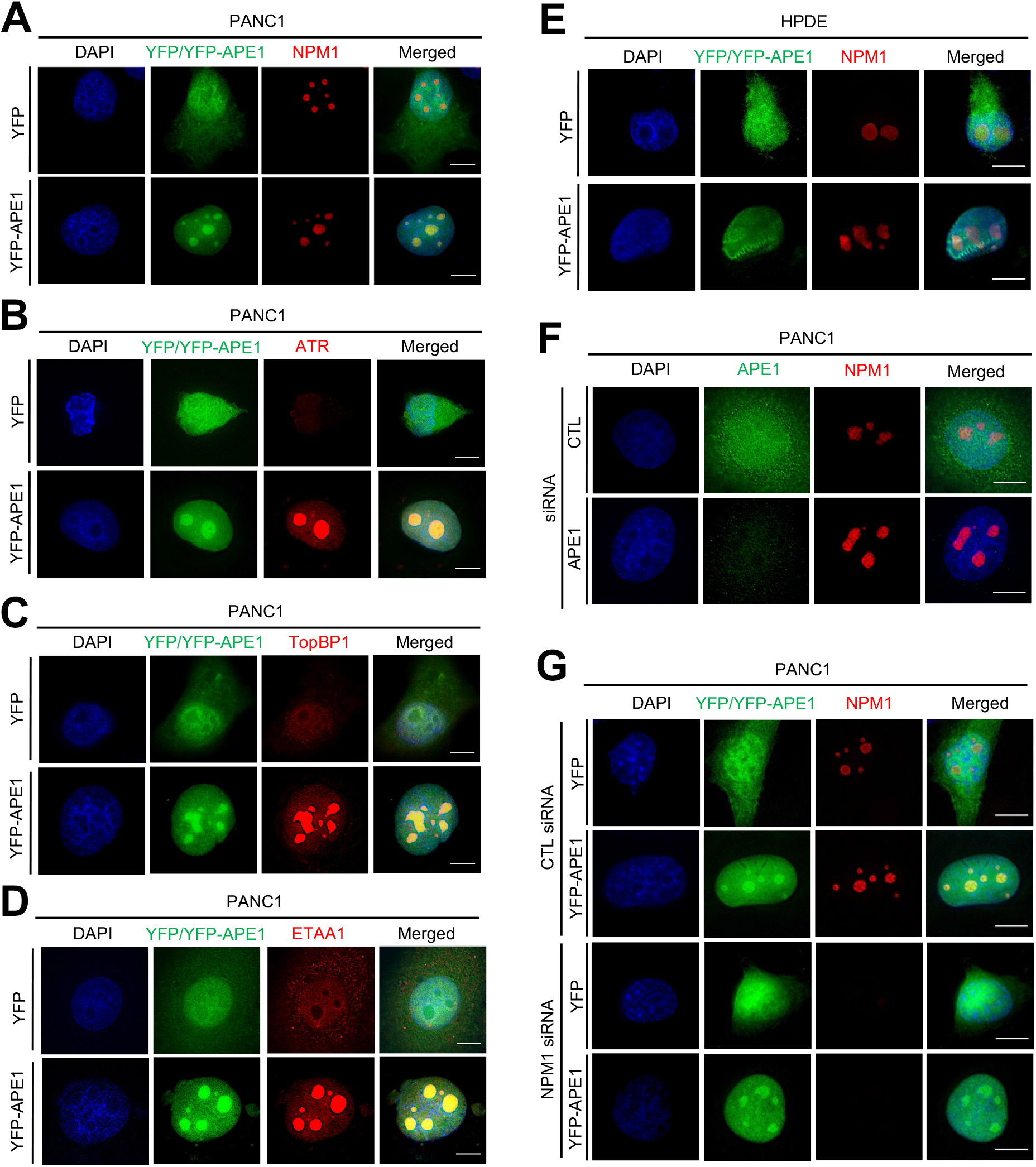
APE1 forms biomolecular condensates in nucleoli to recruit ATR, TopBP1, and ETAA1 for ATR activation in a NPM1-independent fashion in cancer cells but unmalignant cells. (**A**) Overexpressed YFP-APE1 but not YFP colocalized with NPM1 in nucleoli in PANC1 cells. After overexpression of YFP or YFP-APE1 for 3 days, PANC1 cells were fixed and incubated with anti-NPM1-AF647 fluorescence antibody for overnight at 4°C. Then the cells were examined via fluorescence microscopy analysis. (**B-D**) ATR and its activators TopBP1 and ETAA1 were colocalized with YFP-APE1 in nucleoli in PANC1 cells. YFP or YFP-APE1 over-expression plasmid was added to PANC1 cells. After overexpression of YFP or YFP-APE1 for 3d, PANC1 cells were fixed and incubated with anti-ATR-AF647 fluorescence antibody (**B**), anti-TopBP1-AF647 fluorescence antibody (**C**), or anti-ETAA1-AF647 fluorescence antibody (**D**) for overnight at 4°C. Then the cells were analyzed via fluorescence microscopy analysis. (**E**) Overexpressed YFP-APE1 was not colocalized with NPM1 in nucleoli in HPDE cells. Similar experiment to Panel (a) was performed in HPDE cells. Scale bar = 10 μm. (**F**) Nucleolar localization of NPM1 was not affected when endogenous APE1 was knocked down via siRNA. Control (CTL) siRNA or APE1 siRNA was added and transfected to PANC1 cells for 3 days. The cells were then fixed and incubated with anti-APE1-AF488 and anti-NPM1-AF647 fluorescence antibodies for overnight at 4°C, followed by fluorescence microscopy analysis. (**G**) YFP-APE1 still assembled condensates when endogenous NPM1 was knocked down. CTL siRNA or NPM1 siRNA was transfected to PANC1 cells and incubated for 1 day. Then YFP or YFP-APE1 overexpression plasmid was transfected and incubated for 3 days. The cells were fixed and incubated with anti-NPM1-AF647 fluorescence antibody for overnight at 4°C, followed by fluorescence microscopy analysis. Scale bars = 10 μm.

Due to the previously characterized interaction between APE1 and NPM1 within nucleoli in HeLa cells (53), we sought to determine the dependency of APE1 and NPM1 to nucleoli in PANC1 cells. We found that after endogenous APE1 was knocked down via siRNA, NPM1 expression was not affected and NPM1 was still localized within nucleoli (Figure 5F and Supplementary Figure S5D). Similarly, YFP-APE1 still assembled condensates, when endogenous NPM1 was knocked down (Figure 5G and Supplementary Figure S5E). NPM1-knowckdown had almost no noticeable effect on the APE1-OE-induced Chk1 phosphorylation (Supplementary Figure S5E). These observations suggest that NPM1 is dispensable for the nucleolar condensation of YFP-APE1 and associated ATR nDDR.

Furthermore, endogenous APE1 was not colocalized with NPM1 in nucleoli regardless of hydrogen peroxide treatment in PANC1 cells (Supplementary Figure S5F). Oxidative DNA damage triggered partial colocalization of endogenous APE1 with ATR, TopBP1, and ETAA1 (Supplementary Figure S5H-I). Considering the pattern of endogenous APE1 complex with ATR, TopBP1, and ETAA1 in oxidative stress, our results suggest that YFP-APE1-OE-induced nDDR is distinct from oxidative stress-induced global ATR-Chk1 DDR pathway.

### APE1 assembles biomolecular condensates in a TopBP1-independent manner in nucleoli to directly activate ATR-Chk1 nDDR in cells under unperturbed conditions

Previous studies have characterized biomolecular condensates through TopBP1 overexpression within nucleoli and LLPS formation by TopBP1 for ATR DDR pathway (43,65,66). Here we tested whether TopBP1 is important for nucleolar localization of YFP-APE1 and subsequent ATR-Chk1 nDDR. Although TopBP1 was recruited to nucleoli when YFP-APE1 but not YFP was expressed under unperturbed conditions, siRNA-mediated TopBP1-KD had no noticeable change for the nucleolar localization of YFP-APE1 (Figure 6A-B). Moreover, the YFP-APE1-induced Chk1 phosphorylation was reduced mildly when TopBP1 was knocked down (Figure 6B). These data suggest that TopBP1 is dispensable for APE1 condensate in nucleoli and that TopBP1 may function downstream of APE1 for partial contribution to ATR-mediated Chk1 phosphorylation.

**Figure 6.**
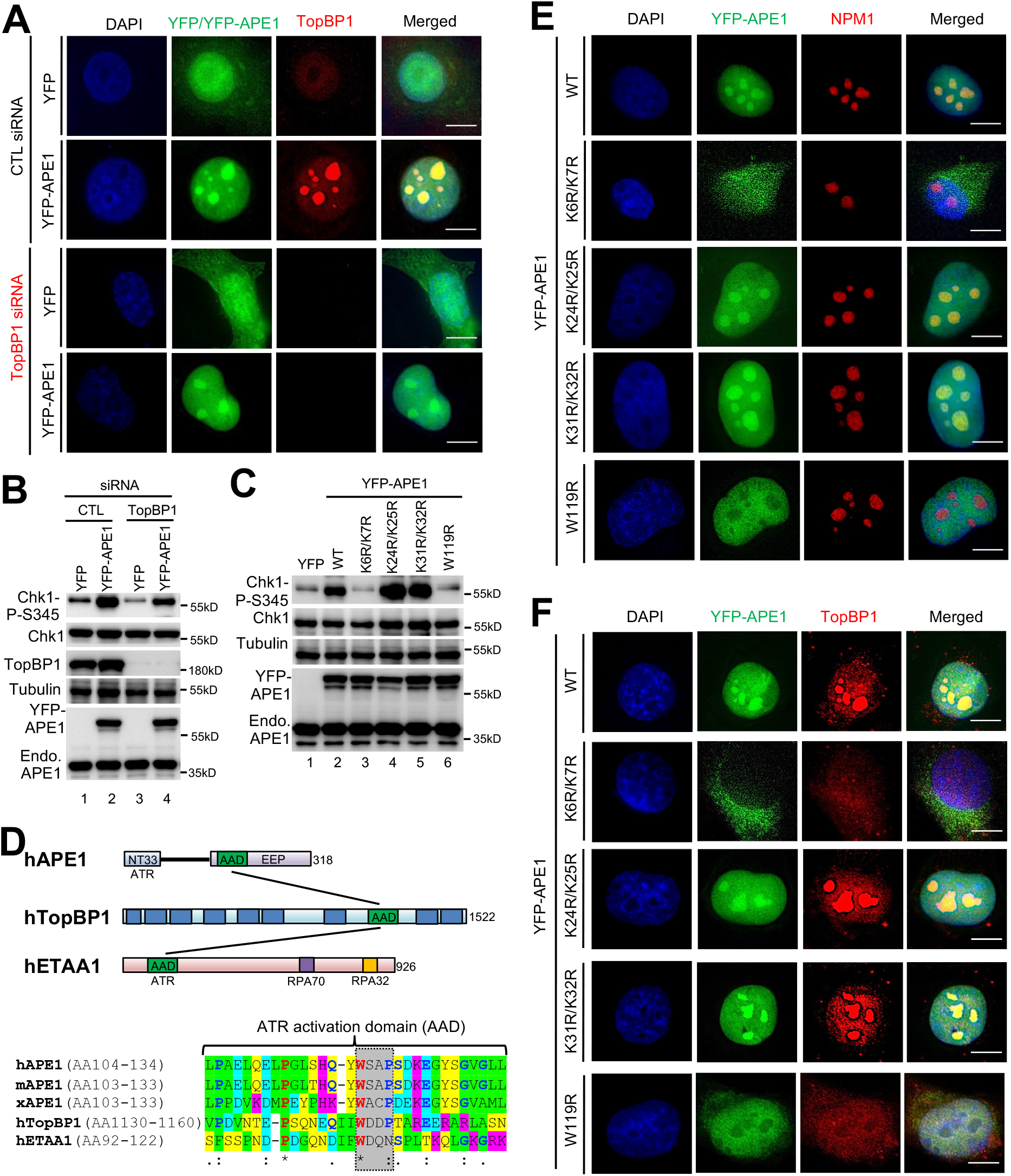
While TopBP1 is dispensable for YFP-APE1 nucleolar localization, the W119 residue within APE1’s putative ATR activation domain is required for its biomolecular condensate formation in nucleoli and nucleolar ATR activation. (**A-B**) YFP-APE1 still assembled condensates and activated ATR-chk1 pathway when endogenous TopBP1 was knocked down via siRNA. After 1-day incubation of CTL or TopBP1 siRNA in PANC1 cells, YFP or YFP-APE1 overexpression plasmid was added and incubation for another 3 days. The cells were fixed and incubated with anti-TopBP1-AF647 fluorescence antibody for fluorescence microscopy analysis (**A**), or extracted for total cell lysates for immunoblotting analysis as indicated (**B**). Scale bar = 10 μm. (**C**) The K6, K7, and W119 residues within APE1 are important for Chk1 phosphorylation induced by YFP-APE1 in PANC1 cells. After YFP, WT or mutant (i.e., K6R/K7R, K24R/K25R, K31R/K32R, W119R) YFP-APE1 overexpression plasmid was added to PANC1 cells and incubation for 3 days, total cell lysates were extracted and analyzed via immunoblotting analysis as indicated. (**D**) Schematic diagrams of functional domains of hAPE1, hTopBP1, and hETAA1 as well as sequence alignment of the putative ATR activation domain (AAD). hAPE1 (human APE1, NCBI#: P27695.1), mAPE1 (mouse APE1, NCBI#: NP_033817.1), xAPE1 (Xenopus laevis APE1, NCBI#: AAH72056.1), hTopBP1 (human TopBP1, NCBI#: Q92547.3), and hETAA1 (human ETAA1, NCBI#: NP_061875.2). *, identical; :, highly conserved; ., low conservation. (**E-F**) APE1 forms biomolecular condensates in nucleoli dependent on its W119 residue. After WT or mutant (i.e., K6R/K7R, K24R/K25R, K31R/K32R, W119R) YFP-APE1 overexpression plasmid was added to PANC1 cells and incubation for 3 days, the cells were fixed and incubated with anti-NPM1-AF647 fluorescence antibody (**E**) or anti-TopBP1-AF647 fluorescence antibody (**F**) for overnight at 4°C, followed by fluorescence microscopy analysis. Scale bar = 10 μm.

Next, we sought to elucidate the molecular determinants within APE1 for nucleolar translocation and LLPS. We first focused on the NT33 motif within APE1 which contains eight conserved Lysine residues (Supplementary Figure S6A). It is previously established that the K6, K7, K24, K25, K27, K31, and K32 within APE1 can be modified by post-translational modifications (PTMs) such as acetylation and/or ubiquitination (80, 81). The conserved K24, K25, K27, K31, and K32 residues are important for interaction with RNA and NPM1 (79, 82). We constructed three double KR mutations (i.e., K6R/K7R, K24R/K25R, and K31R/K32R), in which such mutations remain positively charged but lose accompanying PTMs. WT, K24R/K25R, and K31R/K32R YFP-APE1, but not YFP nor K6R/K7R YFP-APE1 induced Chk1 phosphorylation in PANC1 cells (Figure 6C). Furthermore, K6R/K7R YFP-APE1 was defective in nuclear translocation as well as TopBP1 and ATR colocalization (Figure 6E-F). In contrast, K24R/K25R YFP-APE1 and K31R/K32R YFP-APE1 assembled condensates together with NPM1 within nucleoli and recruited TopBP1 and ATR in a similar fashion to WT YFP-APE1 (Figure 6E-F and Supplementary Figure S6B).

Previous studies on the AAD motifs of TopBP1 and ETAA have shown that the W1145 residue in human TopBP1 (homologue to Xenopus laevis TopBP1 W1138) and the W107 residue in human ETAA1 are critical for ATR activation^23, 24, 81^. Our protein sequence alignment shows that the W119 residue and its nearby ∼30-amino-acid peptide in human APE1 is highly conserved (humans, mouse, and *Xenopus laevis*), and is conserved to some extent to hTopBP1 and hETAA1 (Figure 6D). Notably, W119R YFP-APE1 failed to trigger Chk1 phosphorylation in PANC1 cells (Figure 6C). Although W119R YFP-APE1 was translocated into the nucleus, it was not co-localized with NPM1 and could not form biomolecular condensates within the nucleolus (Figure 6E). Consistent with its defective nucleolar localization, neither TopBP1 nor ATR was recruited to the nucleolus by W119R YFP-APE1 (Figure 6F and Supplementary Figure S6B). These results suggest that the W119 residue within APE1 is important for APE1 LLPS formation into nucleoli *in vivo*, subsequent recruitment of ATR and TopBP1, and ATR-Chk1 nDDR activation. To further test whether the W119 residue is important for APE1’s LLPS *in vitro*, we added WT or W119R His-APE1-YFP to nuclear extracts and found that WT but not W119R His-APE1-YFP formed biomolecular condensates in nuclear extracts (Supplementary Figure S6C). Our *in vitro* pulldown assays from nuclear extracts demonstrated no noticeable change in the association of W119R His-APE1-YFP with ATR, TopBP1, and ETAA1 in comparison to WT His-APE1-YFP (Supplementary Figure S6D).

### APE1-induced nDDR leads to rRNA transcription suppression and impaired cell viability

Based on previous study showing rRNA transcription suppression by TopBP1 overexpression-induced nDDR (43), we sought to determine the potential effect of APE1-induced nDDR in rRNA transcription and cell viability. qRT-PCR assays showed that YPF-APE1 transfection led to reduced pre-rRNA transcription compared with YFP transfection, and such rRNA suppression was sensitive to ATR inhibitor VE-822 (Figure 7A). Moreover, overexpression of K6R/K7R or W119R YFP-APE1 was defective for the rRNA transcription suppression, suggesting that APE1 nuclear and nucleolar localization and LLPS formation are important for rRNA transcription suppression (Figure 7B). These results underscore the significance of APE1-OE-induced nDDR in rRNA transcription suppression.

**Figure 7.**
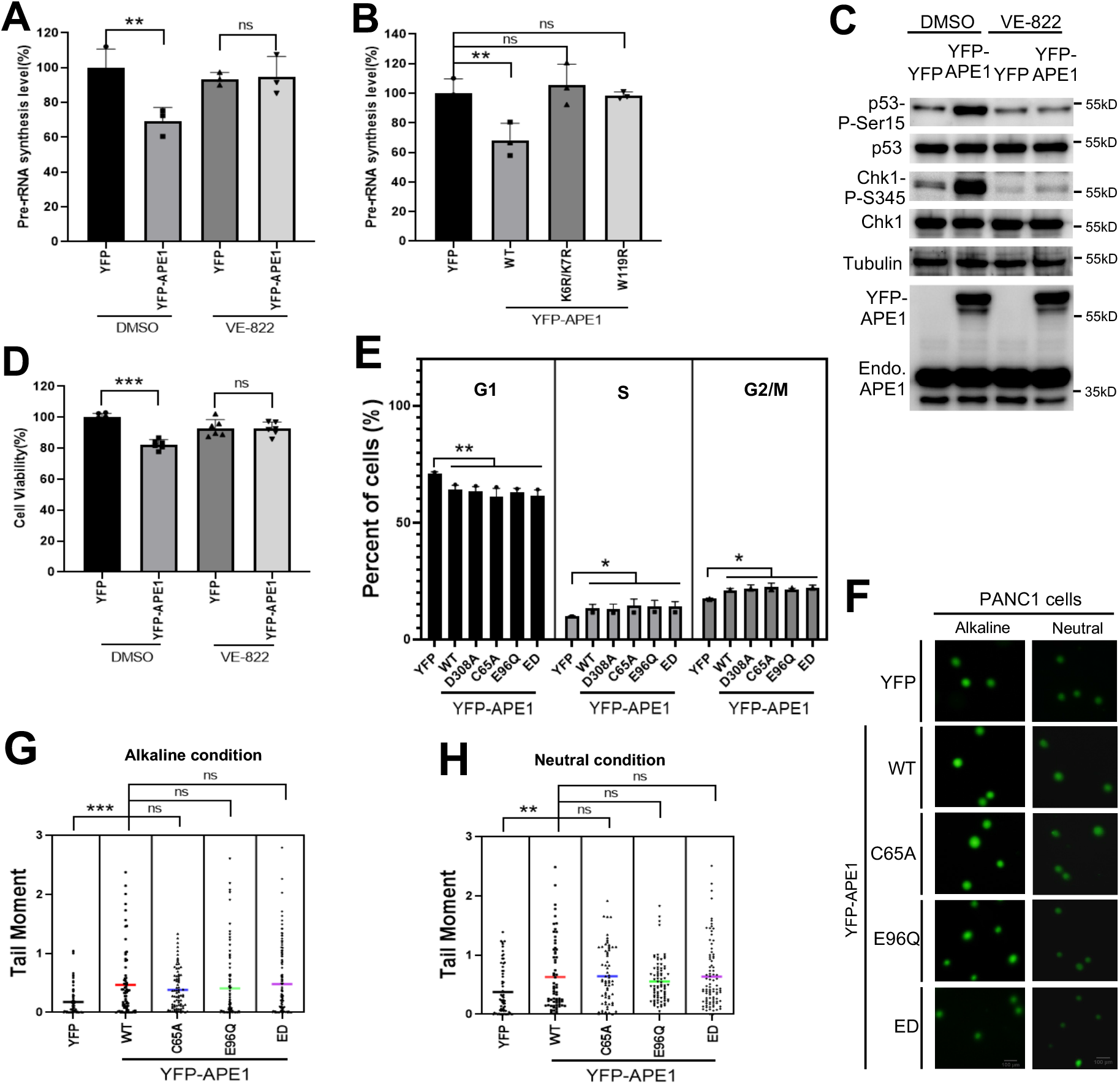
APE1-induced ATR-Chk1 nDDR leads to rRNA transcription suppression and cell viability reduction. (**A**) APE1 overexpression led to pre-rRNA transcription suppression in an ATR-dependent fashion. YFP or YFP-APE1 overexpression plasmid was added to PANC1 cells and incubated for 2 days. DMSO or VE-822 (1μM) was then added and incubated for another day. The total RNA was extracted and analyzed via qRT-PCR, and pre-rRNA synthesis was normalized to beta-Actin. n=3. (**B**) WT but not K6R/K7R nor W119R YFP-APE1 suppressed rRNA transcription. YFP or WT or mutant YFP-APE1 overexpression plasmid was added to PANC1 cells and incubated for 3 days, followed by total RNA extraction and qRT-PCR analysis. n=3. (**C**) Overexpression of YFP-APE1 but not YFP led to p53 Ser15 phosphorylation, which was sensitive to VE-822. After YFP or YFP-APE1 overexpression plasmid was added to PANC1 cells and incubated for 3 days, DMSO or VE-822 was added and incubated for another 2 hours, followed by total cell lysate extraction and immunoblotting analysis as indicated. (**D**) YFP-APE1 overexpression led to cell viability reduction. After YFP or YFP-APE1 overexpression plasmid was added to PANC1 cells and incubated for 2 days, DMSO or VE-822 was added and incubated for another day. The cells were analyzed via MTT assays. n=6. (**E**) Overexpression of WT/mutant APE1 led to cell cycle arrests at S and G2/M phases in PANC1 cells. YFP or WT/mutant YFP-APE1 overexpression plasmid was added to PANC1 cells. After incubation for 4 days, the cells were collected and examined via FACS analysis. The data are presented as means ± SD, n=3. (**F-H**) Overexpression APE1 led to more endogenous DNA damage in PANC1 cells. YFP or WT/mutant YFP-APE1 overexpression plasmid was added to PANC1 cells. After incubation for 4 days, the cells were collected and analyzed via comet assays under alkaline or neutral conditions. Scale bar = 100μm.The data are presented as Scatter dot plot and the lines indicate Means, n>50.

Previous study shows that p53 with a R273H mutation is phosphorylated at Ser15 by gemcitabine-induced stalled DNA replication forks in PANC1 cells (83). To determine the p53 phosphorylation status by APE1-induced nDDR, we found that overexpression of YFP-APE1 but not YFP led to p53 Ser15 phosphorylation, which was sensitive to VE-822 (Figure 7C). We speculated that cell viability might be affected by nDDR-dependent Chk1 and p53 phosphorylation. As expected, our cell viability assays showed that YFP-APE1 overexpression led to about 20% reduction in cell viability compared with YFP transfection and that such cell viability reduction by APE1-OE was reversed by the addition of VE-822 (Figure 7D). These observations suggest the significance of APE1-OE-induced nDDR in p53-mediated apoptosis and cell viability.

Next, we examined cell cycle profiling after overexpression of WT/mutant YFP-APE1. FACS analysis showed that APE1 overexpression led to a higher percentage of PANC1 cells in S phase and G2/M phase regardless of WT or mutant APE1 (Figure 7E). This observation suggests that APE1 overexpression may activate S or G2/M checkpoint responses, consistent to the ATR nDDR pathway activation. Moreover, we analyzed endogenous DNA damage using comet assays after WT/mutant YFP-APE1 overexpression in PANC1 cells. Comet assays showed enhanced tail moment after overexpression of WT and mutant YFP-APE1 in both alkaline condition and neutral condition, compared with YFP (Figure 7F-H), suggesting that endogenous DNA damage (i.e., SSBs, AP sites and DSBs) is upregulated following APE1 overexpression. Overall, APE1 overexpression induces more endogenous DNA damage and S/G2 checkpoint response, which is consistent with the activation of ATR-Chk1 nDDR pathway activation but independent of APE1 nuclease activity and redox function.

Results from this study provide evidence showing that APE1 is important for the global regulation of the ATR-Chk1 DDR pathway under stress conditions via its nuclease activity. It is also shown that overexpressed APE1 forms biomolecular condensates within nucleoli to activate the ATR-Chk1 nDDR pathway. We propose that APE1 contributes to the genome integrity maintenance via distinct regulatory mechanisms under stressful and unperturbed conditions.

## DISCUSSION

### Distinct regulatory mechanisms of APE1 in the ATR-Chk1 DDR pathways

Results from this study provide direct evidence that APE1 contributes to the ATR-Chk1 DDR pathway via distinct molecular mechanisms. Under stress conditions (such as H_2_O_2_, MMS, CPT), stress-induced DNA damage such as AP sites can be recognized and processed by APE1 via its AP endonuclease activity, generating SSB structures with subsequent resection into small gap with 1-3nt by APE1’s 3’-5’ exonuclease activity (Figure 8A). These small ssDNA gaps are further processed by other enzymes such as APE2 to enlarge the RPA-coated ssDNA for ATR/ATRIP recruitment and activation (Figure 8A) (29, 56). Under unperturbed conditions, APE1 overexpression leads to the formation of biomolecular condensates in nucleolus *in vivo* and in nuclear extracts *in vitro* via APE1’s intermolecular and/or intramolecular associations (Figure 8B). Importantly, ATR and its activators TopBP1 and ETAA1 are recruited to APE1-mediated LLPS to promote ATR activation which leads to rRNA transcription suppression in the nucleolus, cell cycle arrest, elevated DNA damage, and reduced cell viability (Figure 8B). To the best of our knowledge, this is the first evidence that APE1 forms LLPS *in vivo* and *in vitro* leading to ATR-Chk1 DDR pathway activation. Intriguingly, such APE1-dependent LLPS-mediated ATR DDR activation in nucleoli is independent of APE1 nuclease and redox function. Thereby, here we define a previously uncharacterized but significant non-catalytic function of APE1 in the ATR-Chk1 DDR pathway.

**Figure 8.**
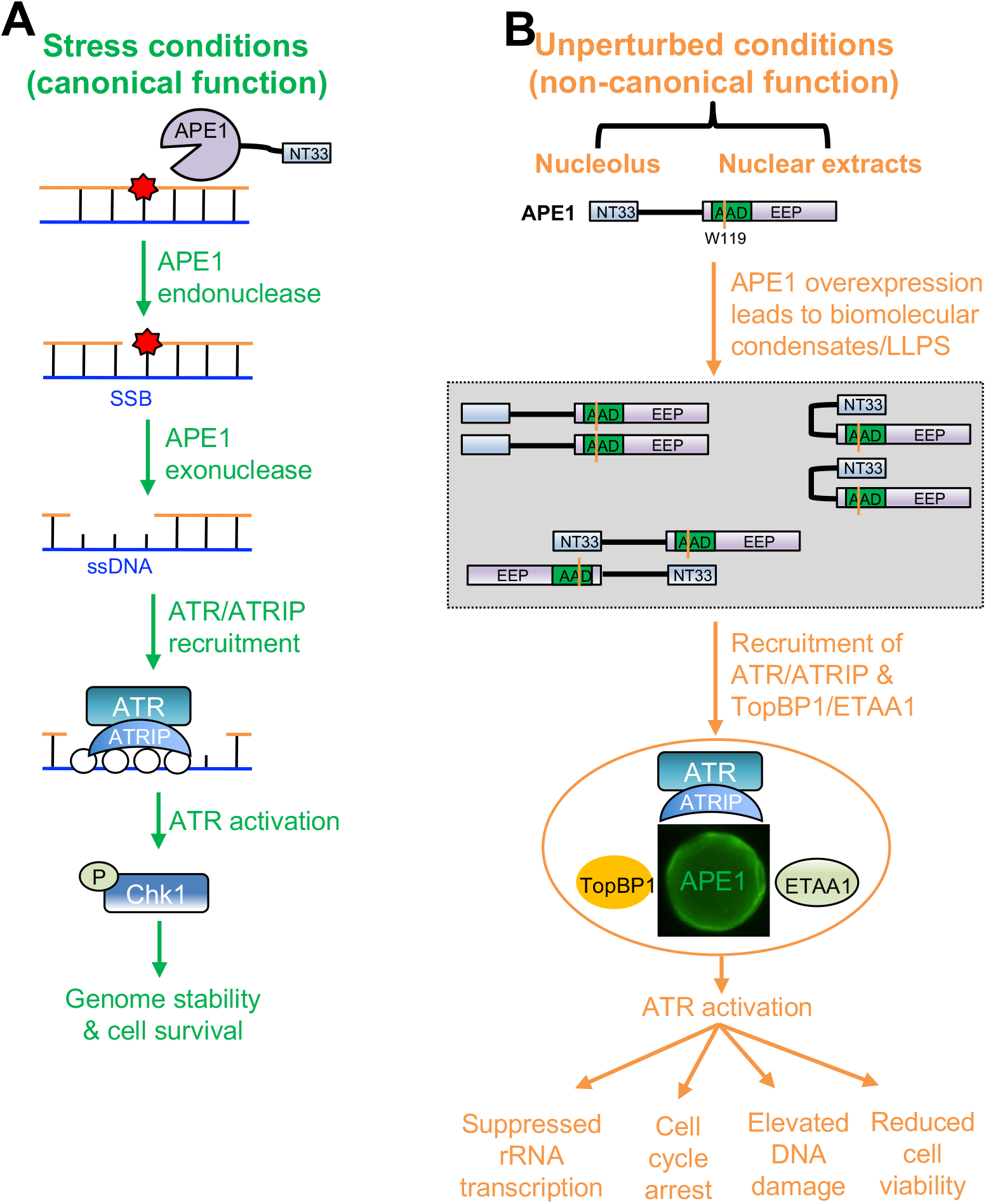
A working model of distinct mechanisms of APE1 in the ATR DNA damage response. (**A**) Stress conditions (canonical function): APE1 promotes the ATR-Chk1 DDR pathway activation via its AP endonuclease and exonuclease activities under stress conditions, which is considered as canonical function of APE1. (**B**) Unperturbed conditions (non-canonical function): under unperturbed conditions, APE1 overexpression leads to biomolecular condensates (liquid-liquid phase separation) in nucleoli *in vivo* and nuclear extracts in vitro via its N33 motif and/or W119 residue in AAD domain. ATR/ATRIP, TopBP1, and ETAA1 are recruited to the APE1-induced condensates for ATR activation. APE1-induced nucleolar ATR DDR suppresses rRNA transcription, arrests cell cycle, elevates DNA damage, and reduces cell viability. See text for more details.

### Canonical and non-canonical function of APE1 in ATR activation

It is recently demonstrated that ATR-Chk1 DDR pathway activation in response to defined SSB structures requires APE1, especially its exonuclease activity, in *Xenopus* egg extracts (56). Such requirement of APE1 and its exonuclease activity in ATR DDR pathway is also conserved in mammalian cells (Figure 1 and Supplementary Figure S1). Thus, APE1’s nuclease activity is considered a canonical function in the ATR DDR. Moreover, our data here demonstrate nucleolar ATR-Chk1 DDR pathway induction via YFP-APE1 that is independent of APE1 nuclease and redox functions. This newly defined YFP-APE1-induced nDDR is dependent on the LLPS assembled by APE1 NT33 motif. This new function of APE1 LLPS in nDDR is here proposed as a non-canonical function.

It is widely accepted that TopBP1 and ETAA1 are direct activators of ATR kinase (23,24,84,85). Here we show that over-expression of APE1 in MDA-MB-231 cells and PANC1 cells can directly activate ATR, independent of APE1 nuclease activity and redox activity (Figure 2). We also provide evidence that recombinant APE1 protein interacts with ATR, ATRIP, RPA, TopBP1 and ETAA to directly activate ATR pathway in *in vitro* nuclear extracts in a NT33 motif-dependent but nuclease- and redox-independent manner (Figures 3-4). Moreover, we identify an ATR activation domain (AAD) similar to TopBP1 and ETAA1’s AAD domain that is required for the translocation and condensation of YFP-APE1 to nucleoli and recruitment of ATR, TopBP1, and ETAA1 (Figure 6). Intriguingly, W1138R AAD within xTopBP1 (homologue to W1145R in hTopBP1) still associates with ATR but is deficient in triggering ATR kinase (84). In contrast, W107A mutant in ETAA1 AAD is defective for both ATR association and ATR activation (23). Our data show that W119R YFP-APE1 is deficient in triggering nDDR and that W119R mutant within His-APE1-YFP has no noticeable change in association with ATR (Supplementary Figure S6D), which agrees with TopBP1 AAD features. Although future *in vitro* kinase assay is needed to directly test this concept, we propose APE1 as a previously unidentified but significant direct activator of ATR kinase.

### LLPS formation by APE1-mediated biomolecular condensates in vitro and in vivo

The N-terminal motif of APE1 can be modified by PTMs including acetylation, ubiquitination and phosphorylation and has been shown association with NPM1, XRCC1, as well as RNA (55). Our data indicate that ATR, ATRIP, TopBP1, ETAA, and RPA32 can interact with APE1 via its N-terminal motif, although it remains unknown whether these proteins’ interaction with APE1 NT33 is direct or indirect (Figures 3-4). Our observation of the defective nuclear location of K6R/K7R YFP-APE1 suggests a potential role for PTMs such as acetylation in K6 and/or K7 residues in the regulation of APE1’s nuclear localization (Figure 6). Future studies are needed to better understand this in greater detail.

The data shown here is the first evidence of APE1-mediated LLPS formation *in vitro* (Figure 4). Our data support the recently proposed concept of APE1’s involvement in LLPS by Tell and colleagues(67, 68). *In vitro* LLPS by APE1 was not dependent on DNA, RNA, or APE1 nuclease/redox functions (Figure 4). Importantly, NT33 motif of APE1 is indeed required for APE1 LLPS *in vitro* (Figure 4). LLPS is often driven by intrinsically disordered protein regions (IDRs) and/or association with nucleic acids such as RNA (86–88). Consistently, the NT33 motif is APE1’s IDR whose structure has not been characterized yet, although other, more ridged, regions of APE1 have been fully characterized (89, 90). We speculate that the bypassed requirement of APE1 NT33 for LLPS in the presence of DNA/RNA is likely due to a compensatory role of another motif within APE1 (Figure 4I-M). Intriguingly, the W119R mutant within APE1 leads to defective APE1 LLPS in nuclear extracts (Supplementary Figure S6C). However, future in-depth studies are needed to determine whether intramolecular interaction within APE1 between APE1 NT33 and a second motif (e.g., W119-containing motif) or intermolecular interaction of APE1 is important for APE1 LLPS. Nonetheless, the newly defined requirement of APE1 NT33 in the *in vitro* LLPS provides new insights into how APE1’s NT33 motif folds and/or associates with other regions of APE1 or its binding partner proteins.

Although previous studies have shown the nucleolar localization of FLAG-APE1 (53, 82), we propose the nucleolar condensation of YFP-APE1 as APE1-assembled LLPS *in vivo* independent of NPM1 but dependent on APE1 W119 residue (Figures 5-6). We also noticed the size and number of YFP-APE1-dependent LLPS in nucleoli varied from one cell to another, while NPM1 typically formed similar size condensates within the nucleoli when YFP was expressed. Whether rDNA, transcribed pre-rRNA and further processed products, or a combination of these factors are important for the APE1 LLPS *in vivo* remains to be determined. Our data reveal the *in vivo* APE1 LLPS within nucleoli subsequently recruits ATR as well as its activators TopBP1 and ETAA1 (Figure 5).

A previous study has revealed that ectopically overexpressed eGFP-TopBP1 is translocated to nucleoli and co-localized with nucleolar protein UBF and RNA Pol I and that TopBP1 overexpression leads to relocalization of NPM1 and NCL from the center region of nucleoli to the peri-nucleolar region (43). Here we show that the localization of TopBP1 induced by YFP-APE1-OE is across almost all the nucleoli (Figures 5-6). While YFP-APE1 is co-localized with NPM1, APE1-induced LLPS doesn’t change NPM1 localization within nucleoli, demonstrating that APE1 and NPM1 nucleolar localization are independent events (Figures 5-6). Recent study shows that the TopBP1-assembled nuclear LLPS depends on ATR-mediated TopBP1 phosphorylation at Ser1138 which serves as positive feedback to amplify the ATR-Chk1 DDR pathway (65). Our data show YFP-APE1-mediated LLPS within nucleoli is independent of TopBP1, indicating APE1 may serve as an upstream regulator of TopBP1 in the context of nucleolar LLPS (Figure 6).

### Physiological significance of ATR DDR pathway activation in nucleoli

YFP-APE1 colocalization and condensation with NPM1 suggests that APE1 may be present in all three regions of nucleoli (Figures 5-6). YFP-APE1-induced ATR nDDR inhibits pre-rRNA transcription (Figure 7), suggesting that the activated ATR may be located in the FC or at the boundary of FC/DFC within nucleoli. However, we can’t exclude the possibility that ATR is recruited to one specific sub-nucleolar compartment by APE1 LLPS and then re-located to another sub-nucleolar compartment once activated.

TopBP1 overexpression leads to ATR-dependent nucleolar segregation but not cell cycle arrest, as well as p53 phosphorylation but not Chk1 phosphorylation (43). Here we demonstrate that YFP-APE1-OE leads to nucleolar translocation and condensation, ATR nDDR-dependent Chk1 and p53 phosphorylation, and cell cycle arrest (Figs. 5-7). This suggests that APE1-induced nDDR is distinguished from TopBP1-induced nDDR pathway although both share some similar features (e.g., rRNA transcription suppression). In addition, two recent studies have demonstrated the requirement of Treacle and TopBP1 but not ETAA1 in the activation of nucleolar ATR DDR pathway in response to DNA replication stress or defined DSBs in rDNA (40, 42). Of note, the nucleolar ATR DDR activation induced by YFP-APE1-OE in this study is under unperturbed conditions in cancer cells but not in normal cells (Figure 5). Considering overall APE1 up-regulation in cancer patients (73,91,92), the APE1-OE-induced ATR nDDR activation provides a new insight into a previously uncharacterized but significant mechanism of the faulty DDR activation by APE1 overexpression in nucleoli of cancer but not normal cells.

In summary, the results of this study uncover that APE1 plays canonical and non-canonical functions in the regulation of ATR-Chk1 DDR pathway (Figure 8). In particular, our study shows for the first time APE1-induced LLPS *in vitro* and *in vivo* and the requirement of APE1-assembled biomolecular condensates in the nucleolar ATR-Chk1 DDR activation. Taken together, the distinctive molecular mechanisms of APE1 in ATR-Chk1 DDR pathway may provide new avenues for future cancer therapies.

## ACKNOWLEDGEMENT

We thank Dr. Pinku Muhkerjee, Dr. Primo Schaer, Dr. Doug Golenbock for reagents. We are grateful for the assistance by Dr. Didier Dréau, Dr. Christine Richardson, Dr. Paola Lopez-Duarte, Dr. Richard Chi, Trey Grissom, and David Gray for FACS as well as fluorescence microscopy analysis.

## Author contributions

J.L. and S.Y. designed experiments. J.L., H.Z. performed experiments. J.L., H.Z., A.M. and S.Y. analyzed the data. J.L. and S.Y. wrote the manuscript. J.L., H.Z., A.M. and S.Y. revised the manuscript. S.Y. supervised the project.

## FUNDING

National Cancer Institute of the National Institutes of Health [R01CA225637 and R01CA251141-subaward to S.Y.]; National Institutes of Environmental Health Sciences of the National Institutes of Health [R21ES032966 to S.Y.]; University of North Carolina at Charlotte [Internal Funding to S.Y.]. Funding for open access charge: the National Institutes of Health.

## Competing interests

None declared.

## SUPPLEMENTARY DATA

**Supplementary Data** are available….

## Supplementary Data (Supplementary Figures)

**Figure S1.**
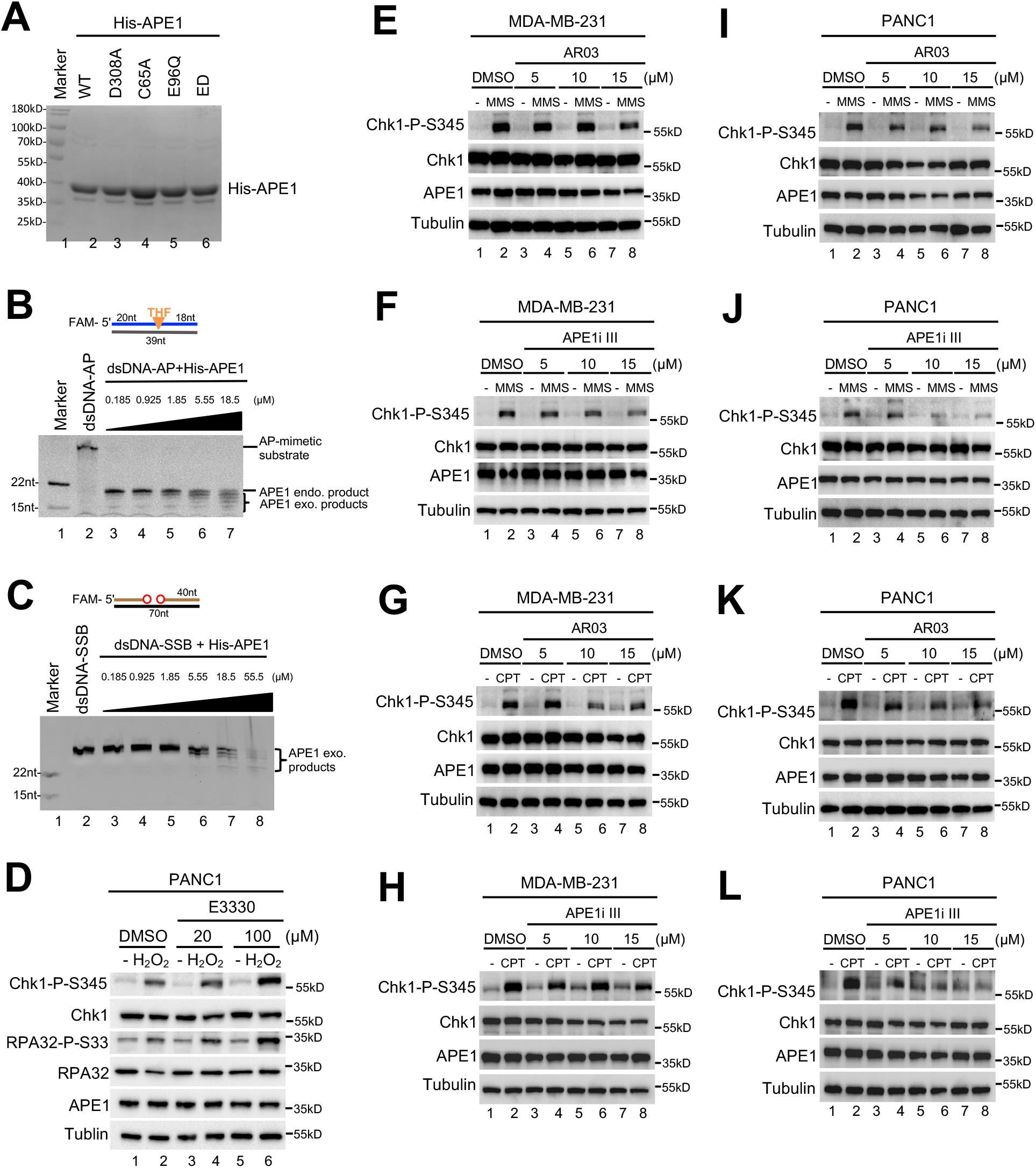
APE1 is important for ATR-Chk1 DDR pathway activation in MDA-MB-231 and PANC1 cells. (**A**) Purified WT or mutant His-APE1 protein was examined on SDS-PAGE gel and stained via coomassie brilliant blue. (**B**) Recombinant His-APE1 protein displayed endonuclease activity. Different doses of WT His-APE1 protein was added to endonuclease assays containing dsDNA-AP as substrate for 30-minute incubation at 37°C, followed by examination via 20% TBE gel. (**C**) Recombinant His-APE1 protein displayed exonuclease activity. Different doses of WT His-APE1 protein was added to endonuclease assays containing dsDNA-SSB as substrate for 30-minute incubation at 37°C, followed by examination via 20% TBE gel. (**D**) E3330 had almost no effect on the H_2_O_2_-induced Chk1 phosphorylation in MDA-MB-231 cells. DMSO or different doses of E3330 as indicated was added to MDA-MB-231 cells for 2 hours, followed by addition of H_2_O_2_ (1.25 mM) for 2 hours. Total cell lysates were examined via immunoblotting analysis as indicated. (**E-H**) AR03 or APE1i III impaired Chk1 phosphorylation induced by MMS or CPT in MDA-MB-231 cells. After different doses of AR03 or APE1i III (5, 10, 15 μM) was added to MDA-MB-231 cells for 2 hours, cells were treated with MMS (0.3 mg/mL, **E-F**) or CPT (10 μM, **G-H**) for 2 hours. Total cell lysates were then examined via immunoblotting analysis as indicated. (**I-L**) AR03 or APE1i III compromised Chk1 phosphorylation induced by MMS or CPT in PANC1 cells. The similar experiments as in (**E-H**) were performed in PANC1 cells.

**Figure S2.**
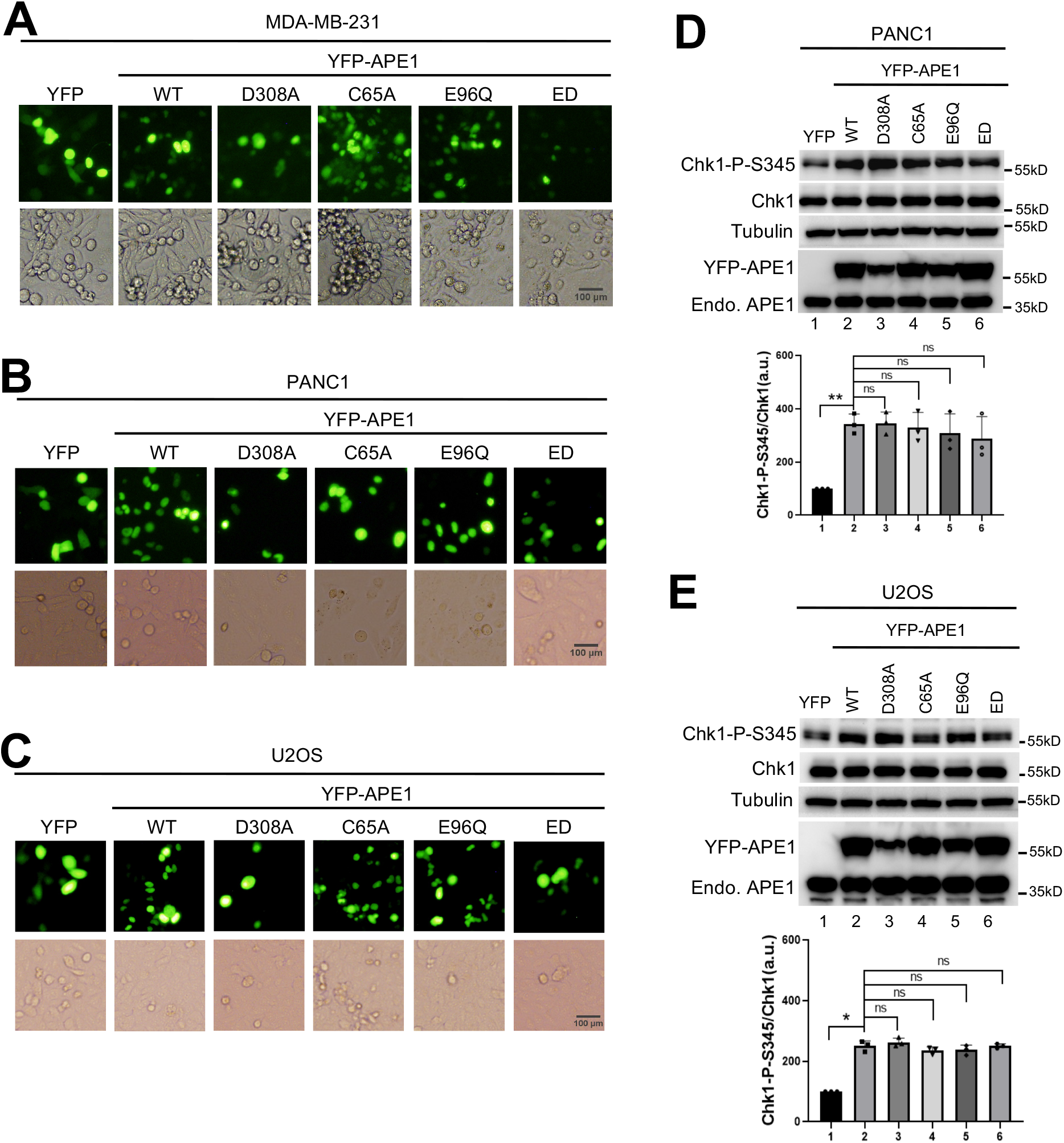
Overexpression of WT/mutant YFP-APE1 triggers Chk1 phosphorylation in different cell lines. (**A-C**) YFP, WT or mutant YFP-APE1 overexpression plasmid was added to MDA-MB-231 (**A**), PANC1 (**B**), or U2OS (**C**) cells. After incubation for 4 days, YFP or YFP-APE1 expression was examined via fluorescence microscopy analysis. Scale bar = 100 μm. (**D-E**) Overexpression of WT/mutant YFP-APE1 protein triggered Chk1 phosphorylation independent of its endonuclease of exonuclease activity in PANC1 or U2OS cells. After 4-day incubation of YFP or WT/mutant YFP-APE1 expression in PANC1 (**D**) or U2OS (**E**) cells, total cell lysates were extracted and analyzed via immunoblotting analysis as indicated. Chk1-P/Chk1 were quantified and presented as means ± SD (n=3).

**Figure S3.**
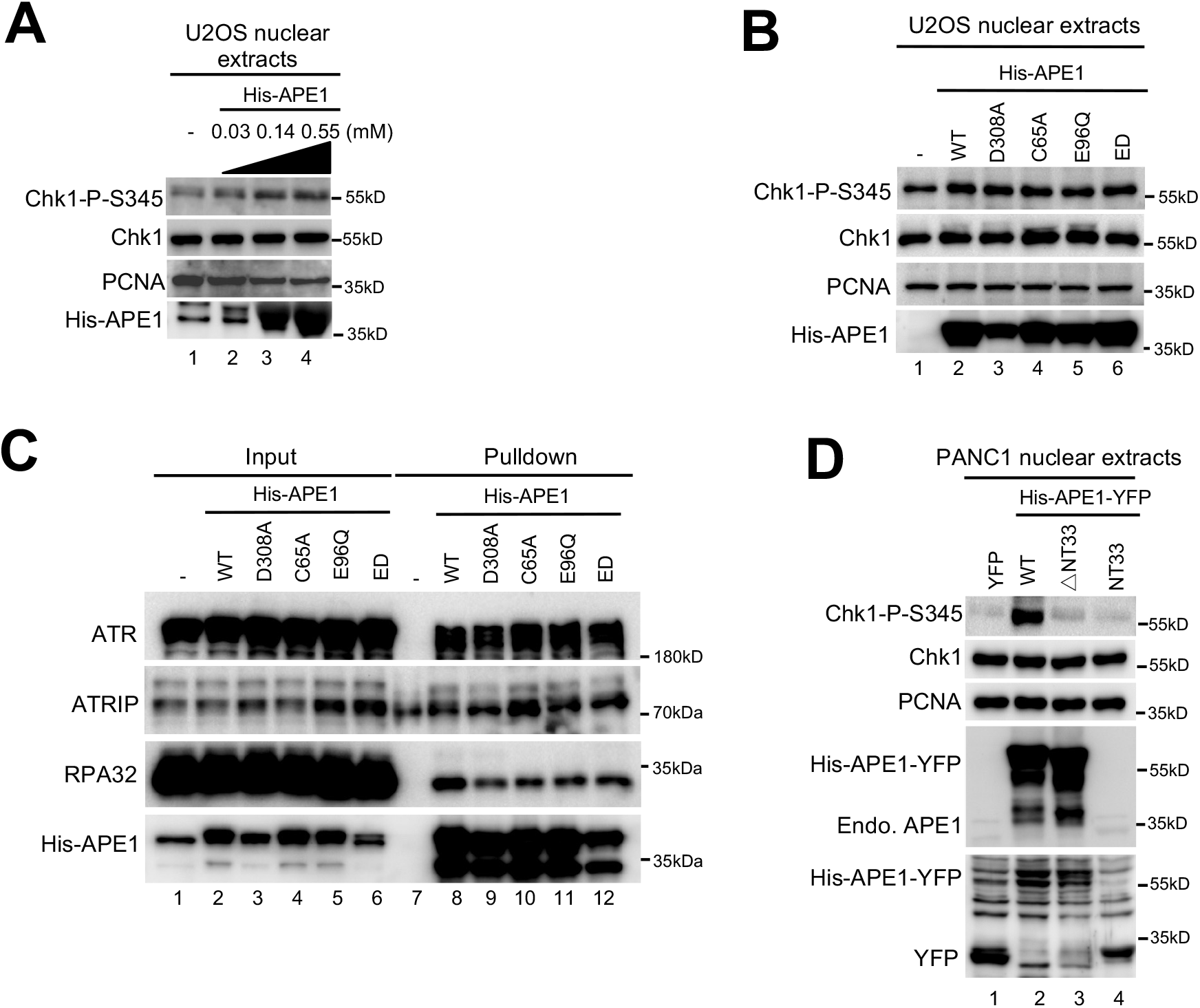
Recombinant APE1 protein activates Chk1 phosphorylation in nuclear extracts from U2OS and PANC1 cells. (**A**) His-APE1 protein activated ATR-Chk1 DDR pathway in a dose-dependent fashion in U2OS nuclear extracts. Different doses recombinant His-APE1 protein was added to U2OS nuclear extracts and incubated for 30 minutes at room temperature. Total cell lysates were examined via immunoblotting analysis as indicated. (**B**) Chk1 phosphorylation induced by His-APE1 protein in U2OS nuclear extracts was independent of it nuclease activity or redox function. After the excess addition of WT/mutant His-APE1 protein to U2OS nuclear extracts and incubation for 30 minutes at room temperature, the samples were analyzed via immunoblotting analysis as indicated. (**C**) Pulldown assays showed that WT/mutant His-APE1 interacted with ATR, ATRIP, and RPA32 in U2OS nuclear extracts. “Input” and “Pulldown” samples were analyzed via immunoblotting analysis as indicated. (**D**) Neither △NT33 His-APE1-YFP protein nor NT33-His-APE1-YFP protein triggered Chk1 phosphorylation in PANC1 nuclear extracts. Purified recombinant YFP, WT His-APE1-YFP, △NT33 His-APE1-YFP, or NT33 His-APE1-YFP protein was added to PANC1 nuclear extracts and incubated at room temperature for 30 minutes. Total samples were then analyzed via immunoblotting analysis as indicated.

**Figure S4.**
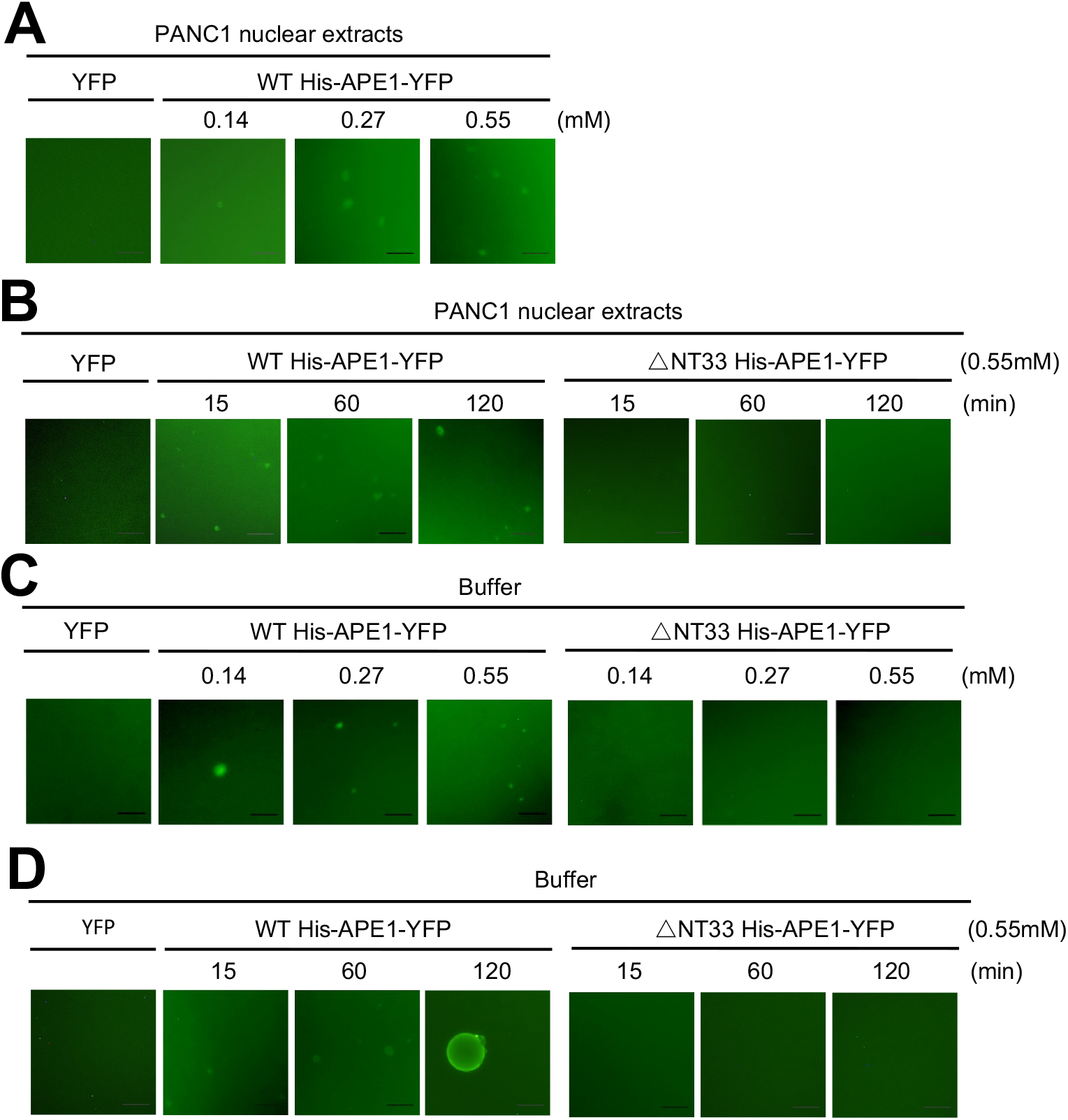
APE1 forms biomolecular condensates in vitro. (**A**) WT His-APE1-YFP formed phase separation in a dose dependent manner in PANC1 nuclear extracts. Different doses of WT His-APE1-YFP (i.e., 0.14, 0.27, or 0.55 mM) or YFP (0.55 mM) was added to PANC1 nuclear extract for 15-minute incubation at room temperature, followed by fluorescence microscopy analysis. Scale bar = 50 μm. (**B**) Time-course experiment of phase separation induced by WT but not △NT33 His-APE1-YFP in PANC1 nuclear extracts. YFP, WT or △NT33 His-APE1-YFP (0.55 mM) was added to nuclear extracts and incubated for different times as indicated. Reaction mixtures were examined by fluorescence microscopy analysis. Scale bar = 50 μm. (**C**) Dose-dependent experiment of phase separation induced by WT but not △NT33 His-APE1-YFP in a LLPS buffer. Different doses of WT or △NT33 His-APE1-YFP (i.e., 0.14, 0.27, or 0.55 mM), or YFP (0.55 mM) was added to a LLPS buffer and incubated for 15 minutes at room temperature, followed by fluorescence microscopy analysis. Scale bar = 50 μm. (**D**) Time-course experiment of phase separation induced by WT but not △NT33 His-APE1-YFP in a LLPS buffer. Similar experimental setup as (**B**) except incubation in a buffer.

**Figure S5.**
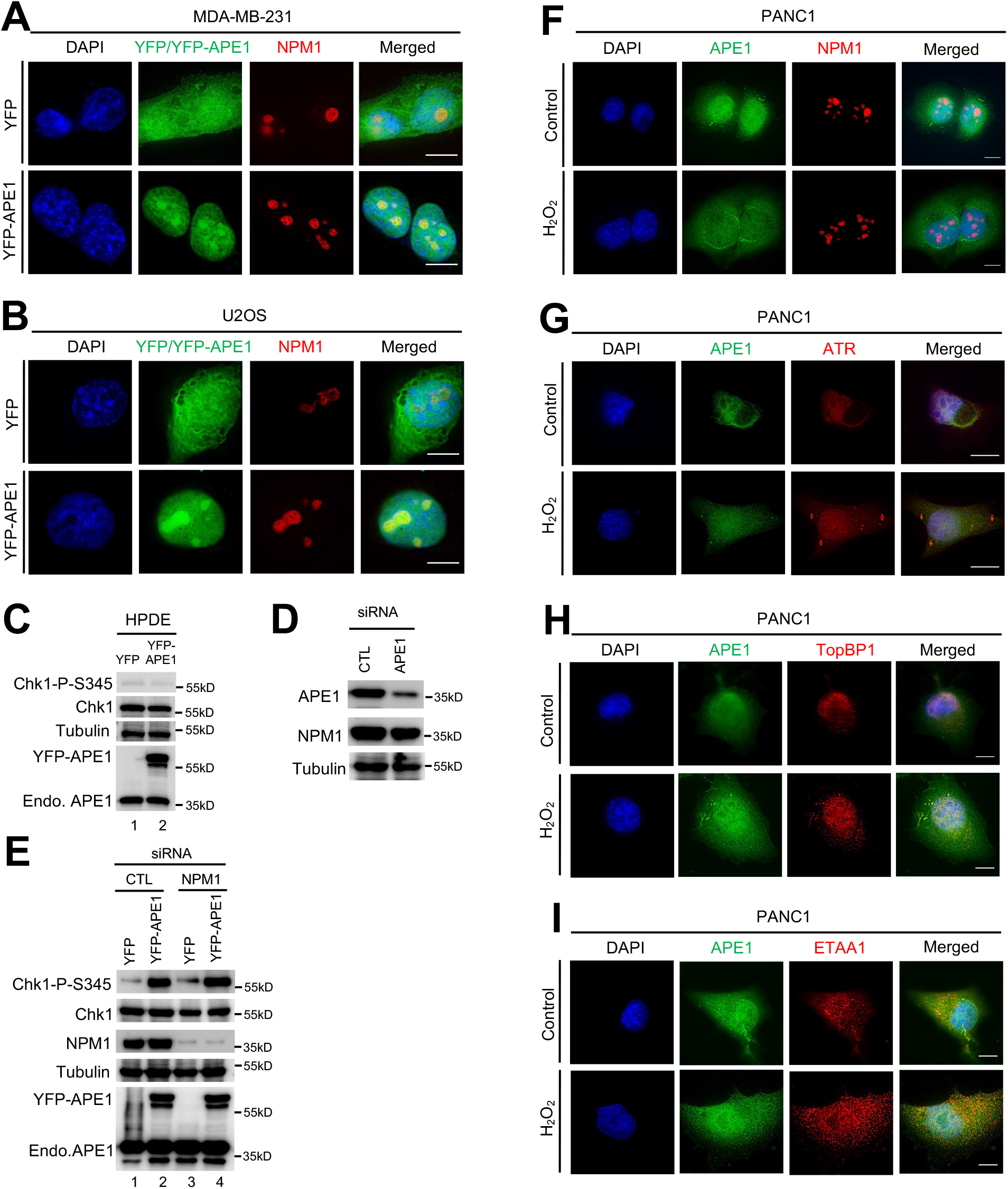
Endogenous APE1 partially colocalizes with ATR, TopBP1 and ETAA1 but not NPM1 in oxidative stress in PANC1 cells. (**A-B**) Overexpressed YFP-APE1 but not YFP colocalized with NPM1 in nucleoli of cancer cells. After overexpression of YFP or YFP-APE1 for 3 days, MDA-MB-231 (**A**) or U2OS (**B**) cells were fixed and incubated with anti-NPM1-AF647 fluorescence antibody for overnight at 4°C. Then the cells were examined via fluorescence microscopy analysis. Scale bar = 10 μm. (**C**) immunoblotting analysis shows Chk1 is not phosphorylated after YFP-APE1 transfection in HPDE cells. (**D**) Endogenous APE1 was reduced via siRNA. Total cell lysates were extracted and analyzed via immunoblotting analysis. (**E**) siRNA-mediated knockdown of endogenous NPM1 had no effect for Chk1 phosphorylation induced by APE1 overexpression. Total cell lysates from panel were extracted and analyzed via immunoblotting analysis as indicated. (**F**) Endogenous APE1 was not colocalized with NPM1 in the nucleoli in PANC1 cells regardless of oxidative stress. H_2_O_2_ (1.25 mM) was added to PANC1 cells for 2h, followed by incubation with anti-APE1-AF488 fluorescence antibody combined with anti-NPM1-AF647 fluorescence antibody for overnight at 4°C. Cells were then analyzed via fluorescence microscopy analysis. Scale bar = 10 μm. (**G-I**) Endogenous APE1 colocalized with ATR, TopBP1 and ETAA1 following oxidative stress in PANC1 cells H_2_O_2_ (1.25 mM) was added to PANC1 cells for 2h, followed by incubation with anti-APE1-AF488 fluorescence antibody in a combination with anti-ATR-AF647 fluorescence antibody (**G**), anti-TopBP1-AF647 fluorescence antibody (**H**), or anti-ETAA1-AF647 fluorescence antibody (**I**) for overnight at 4°C. Cells were then analyzed via fluorescence microscopy analysis. Scale bar = 10 μm.

**Figure S6.**
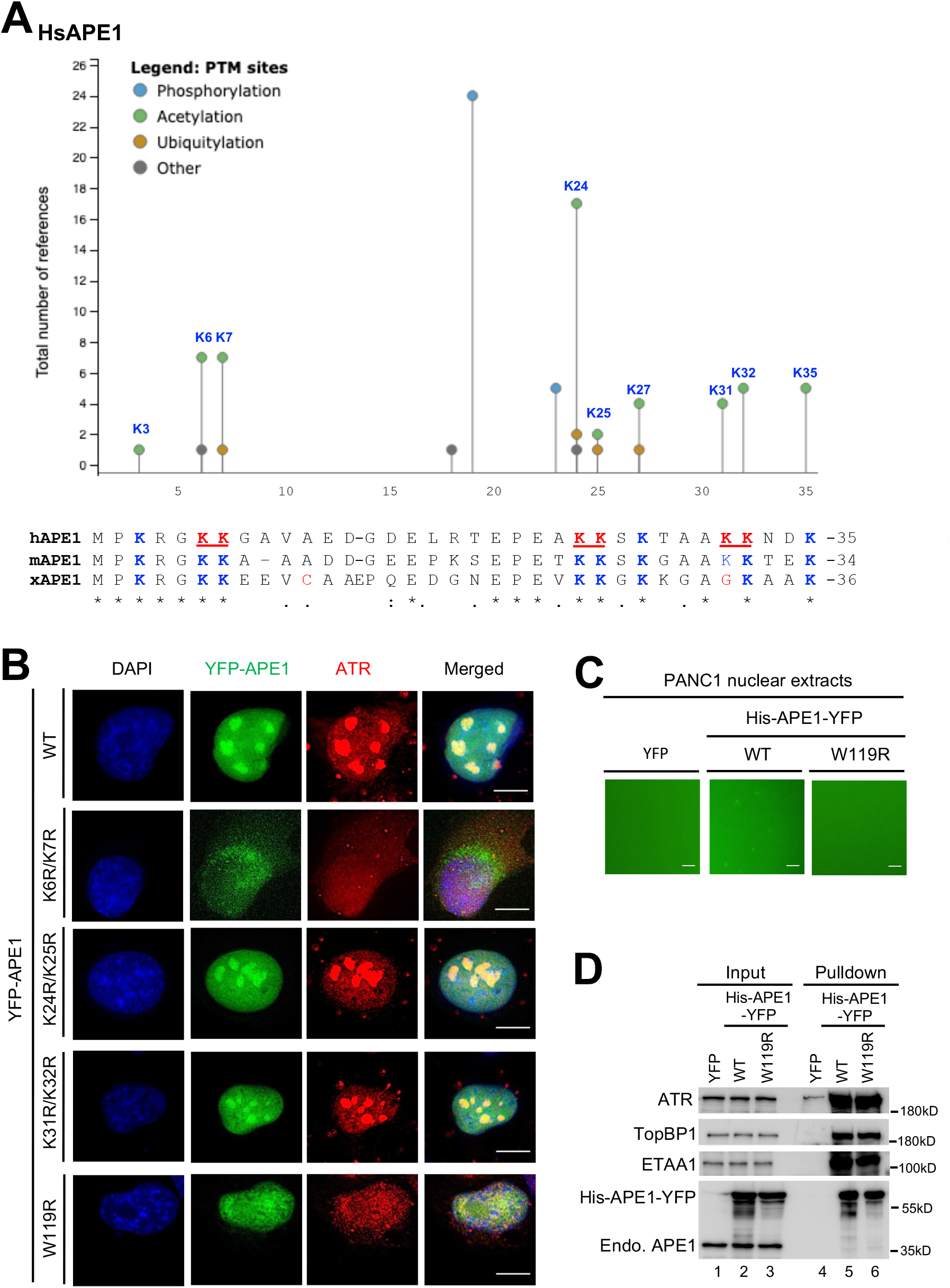
APE1 forms biomolecular condensates in nucleoli dependent on its W119 residue. (**A**) Sequence alignment of the N-terminal motif of hAPE1, mAPE1 and xAPE1. The top diagram shows the PTMs of lysine residues in human APE1 modified from PhosphoSitePlus (www.phosphosite.org) analysis. (**B**) The K6/K7 and W119 residues of YFP-APE1 were required for biomolecular condensates formation within the nucleoli and colocalization of APE1 with ATR. WT or mutant (i.e., K6R/K7R, K24R/K25R, K31R/K32R, W119R) YFP-APE1 overexpression plasmid was transfected into PANC1 cells and incubated for 3 days. The cells were then fixed and incubated with anti-ATR-AF647 fluorescence antibody for overnight at 4°C for fluorescence microscopy analysis. Scale bar = 10μm. (**C**) Biomolecular condensates induced by YFP-APE1 in PANC1 nuclear extracts was dependent on its W119 residue. YFP, WT or △NT33 His-APE1-YFP (0.55 mM) was added to nuclear extracts and incubated for 15 minutes at room temperature, followed by fluorescence microscopy analysis. Scale bar = 50μm. (**D**) W119R His-YFP-APE1 associated with ATR, TopBP1 and ETAA1 from PANC1 nuclear extracts, similar to WT His-YFP-APE1. Input and Pulldown samples were examined via immunoblotting analysis as indicated. (**F**) Chk1 phosphorylation was activated by excess addition of WT His-APE1-YFP protein but not △N33 His-APE1-YFP in PANC1 nuclear extracts. Chk1-P/Chk1 was quantified and analyzed as means ± SD (n=3). (**G**) Chk1 phosphorylation induced by WT His-APE1-YFP protein in PANC1 nuclear extracts was inhibited by VE-822 but not KU55933 nor NU7441 (1 mM). (**H**) Pulldown assays suggest that WT but not △N33 His-APE1-APE1 associated with ATR, ATRIP and RPA32 in PANC1 nuclear extracts. “Input” or “Pulldown” samples were examined via immunoblotting analysis as indicated.

